# Random crosslinks generate anomalous scaling of dynamic moduli of biomolecular condensates

**DOI:** 10.1101/2025.11.05.686888

**Authors:** Bohan Lyu, Jie Lin

**Affiliations:** Peking-Tsinghua Center for Life Sciences, Peking University, Beijing, China; Center for Quantitative Biology, Peking University, Beijing, China

## Abstract

Biomolecular condensates are viscoelastic, and their mechanical properties are intimately related to their biological functions. However, the connection between microscopic networks formed by intermolecular crosslinks and viscoelasticity is still elusive. Here, we model biomolecular condensates as random crosslinked polymer solutions to elucidate how random connectivity fundamentally alters their viscoelasticity. We decompose the entire condensate into multiple clusters without loops and demonstrate that for clusters with size *n*, their eigenvalue distributions exhibit a power-law scaling *p*_*n*_(*λ*) ∼*λ*^*−*1*/*3^ with a lower cutoff *λ*_min_ ∼*n*^*−*3*/*2^. By integrating all clusters, we show that for the entire condensate, random crosslinks generate abundant slow modes involving multiple linear polymers with a constant eigenvalue distribution. The slow modes cause anomalous linear frequency scaling of the dynamic moduli; in particular, they significantly boost the low-frequency storage modulus relative to uncrosslinked systems. Our model rationalizes the anomalous scaling of the dynamic moduli observed in multiple biomolecular condensates.

## Introduction

Biomolecular condensates, also known as membraneless organelles, play crucial roles in cellular organizations and functions, e.g., by concentrating specific proteins and nucleic acids into dynamic compartments such as P granules [1, 2], stress granules [3, 4], transcriptional condensates [5–7], and many others [8–10]. Notably, many experiments have demonstrated that biomolecular condensates are viscoelastic, exhibiting both liquid-like and solid-like behaviors, which are quantified by the storage modulus *G*^′^ and loss modulus *G*^′′^ [11–15]. Importantly, the material properties of condensates also significantly influence their biological functions and may even lead to diseases [16–22].

Experiments and simulations have highlighted the central role of intermolecular interactions in shaping the mechanical properties, morphologies, and aging behaviors of biomolecular condensates [13, 15, 23–29]. Importantly, biomolecular condensates often involve multivalent proteins, e.g., via intrinsically disordered regions (IDRs) or sticky patches, and may form networks, giving rise to emergent viscoelastic behaviors distinct from those of linear polymers [30–32]. Recent computational studies have also shown that accounting for crosslinks between different proteins is critical for matching numerical and experimental data on the dynamic moduli [33, 34]. Moreover, recent experiments have revealed anomalous linear scalings in the low-frequency regime of storage and loss moduli: *G*^′^(*ω*) ∼*ω* and *G*^′′^(*ω*) ∼*ω* where *ω* is the frequency [11, 12, 34], distinct from the predictions of the Maxwell model [35].

Classical theories of polymer viscoelasticity, such as the Rouse and Zimm models, provide foundational frameworks for linear polymers but fall short of capturing the complex rheology of biomolecular condensates [35–38]. To address complex topological features, the Rouse model has been generalized to arbitrary networks, demonstrating that inter-chain crosslinks fundamentally reshape the viscoelastic response [33, 34, 39]. Notably, recent studies have successfully used this generalized framework to reproduce experimentally measured moduli [33, 34]. Nevertheless, these computational approaches rely on input of network topology from independent simulations. Therefore, a tractable minimal model that explicitly reveals the universal origin of this anomalous scaling remains elusive.

In this work, we propose a minimal model of randomly crosslinked polymer solutions below gelation to describe biomolecular condensates, elucidating how random connectivity profoundly transforms their viscoelastic behaviors. We study the eigenvalue distribution of the connectivity matrix of the condensate, because it is the eigenvalues of the connectivity matrix that determine the condensate’s viscoelastic responses [33, 34]. Since the entire condensate consists of multiple separate clusters below gelation, we first investigate the eigenvalue distributions of clusters of a given size. We show that for clusters with size *n*, the eigenvalue distribution averaged over various topologies exhibits a power-law scaling, *p*_*n*_(*λ*) ∼*λ*^−*α*^ for small eigenvalues *λ* with a lower cutoff *λ*_min_(*n*) ∼*n*^−*β*^, where *α* ≈1*/*3 and *β* ≈3*/*2. Notably, the relationship between a cluster’s viscosity and its mean-square radius of gyration [39] allows us to predict the scaling relation, *αβ* = 1*/*2, in agreement with our numerical calculations.

By combining all clusters in the system, we further reveal that the entire condensate exhibits a constant eigenvalue distribution in the limit of small *λ*, reflecting an abundance of slow collective modes. Remarkably, these slow modes lead to anomalous linear scaling in both the storage and loss moduli (with a logarithmic correction) at low frequency, significantly enhancing the low-frequency storage modulus relative to that of uncrosslinked polymer solutions. Our theoretical predictions are consistent with experimental data from several types of biomolecular condensates [11, 12, 34], suggesting that the slow modes generated by random crosslinking provide a general mechanism for this widely observed viscoelastic signature.

### Random crosslinked polymers in a condensate

We propose an analytically tractable minimal model to study the viscoelasticity of biomolecular condensates. We assume that the condensate is composed of strongly overlapping long linear polymer chains, which is generally valid for the dense phase of polymer solution after phase separation [35], and that proximate chain segments have a finite probability of forming a crosslink, e.g., via sticky amino acids or patches at a protein’s surface [23, 40– 42] (Figure 1a), which persists on the experimental timescale.

**FIG. 1.**
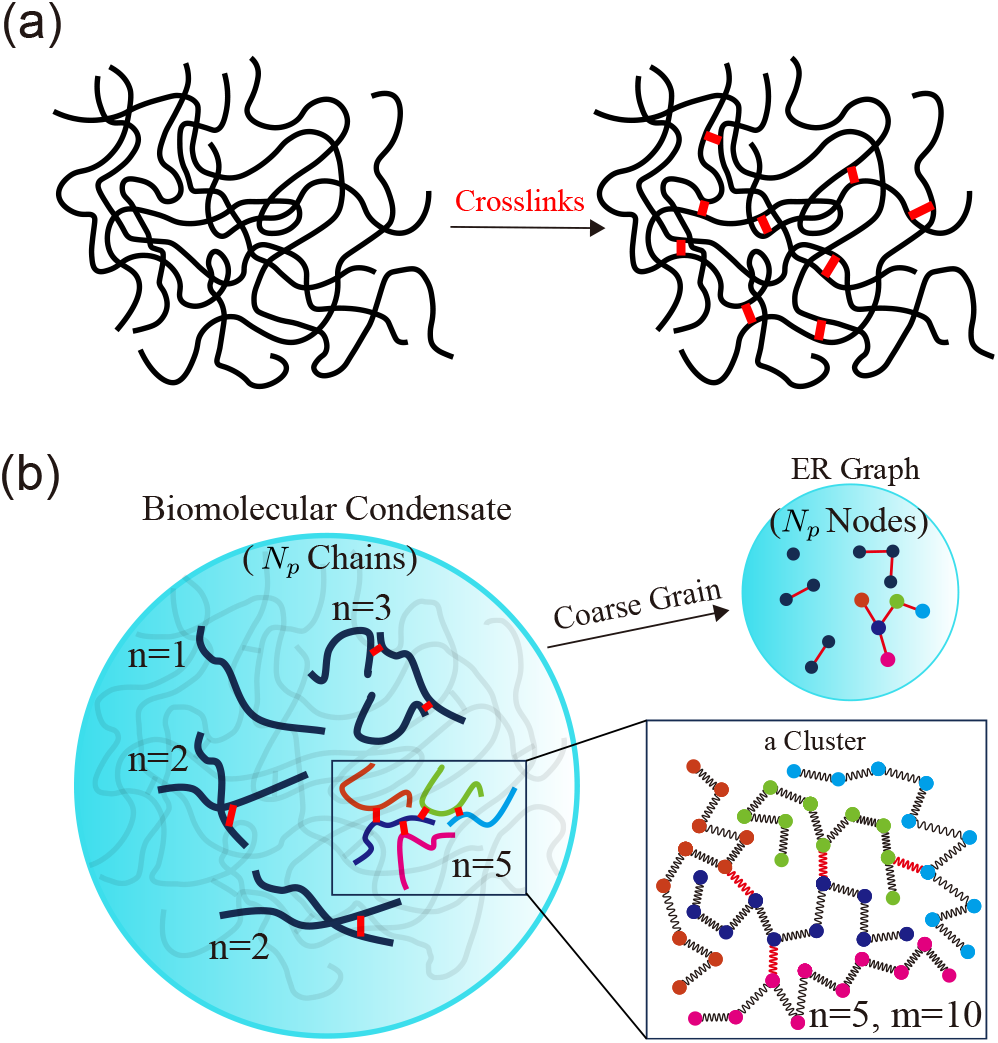
(a) We model biomolecular condensates as a crosslinked polymer solution where linear polymer chains, e.g., proteins, are connected via random crosslinks (red lines). (b) The condensate, containing a total of *N*_*p*_ chains, is composed of multiple clusters of varying sizes, some of which we highlight here. The entire condensate can be coarse-grained into an ER graph by treating each chain as a node and each crosslink as an edge; the coarse-grained graph of the highlighted clusters is shown in the upper-right inset. The lower-right inset shows the structures of a cluster of *n* = 5 chains with 4 crosslinks. Each chain has *m* = 10 beads.

The strong overlap between the long polymer chains justifies the mean-field approximation [35], under which any two chains have an equal probability of being connected by a crosslink, thereby mapping the condensate’s network topology precisely to an Erdős-Rényi (ER) random graph where each chain is coarse-grained as a node (Figure 1b) [43]. We focus on the liquid-like state below the gelation point, where the mean number of crosslinks per chain *c <* 1 (in agreement with the experimental data analyzed later), precluding the formation of an infinite cluster that would otherwise invalidate the mean-field approach. In this subcritical regime, the system naturally decomposes into an ensemble of finite-sized clusters and each cluster is connected without loops (Figure 1b; see the detailed proof in Supplemental Material Section D1).

For a condensate with *N*_*p*_ chains, the number of crosslinks is *N*_*p*_*c/*2. In the thermodynamic limit (*N*_*p*_→ ∞) below gelation threshold (*c <* 1), the probability distribution for a randomly selected chain to belong to a cluster of size *n* (the number of chains in the cluster) is given by (Supplemental Material Section D1)

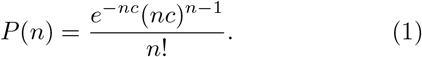

For *n* ≫1, 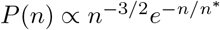, where *n*^*^ ∝ (1 −*c*)^−2^.

### Generalized Rouse model for crosslinked networks

We use the generalized Rouse model where the crosslinked polymer solution is represented by beads (i.e., monomers) connected by springs [33, 34, 39], and each polymer chain has *m* beads (Figure 1b). The position of bead *i* in three dimensions follows overdamped dynamics

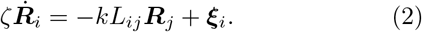

Here, *ζ* is the friction coefficient, which implicitly includes the effects of transient contacts, and *k* is the spring constant, which can be written as *k* = 3*k*_*B*_*T/b*^2^ where *k*_*B*_ is the Boltzmann constant, *T* is the temperature, and *b*^2^ is the mean square distance between two connected beads. For simplicity, we use the same spring constant for polymer backbones and crosslinks, which does not affect our main conclusions. The random force ***ξ***_*i*_ is generated by thermal fluctuation and satisfies the fluctuation-dissipation theorem (FDT) [44]. The matrix *L*_*ij*_ is symmetric and describes the connectivity between beads, where *L*_*ij*_ = −1 if beads *i* and *j* are connected, *L*_*ii*_ is the total number of connections to bead *i*, and otherwise *L*_*ij*_ = 0. This matrix has non-negative eigenvalues *λ*_*p*_ [33, 37, 39]. We note that Eq. (2) can be applied to any networks, either a single cluster or the entire condensate.

The relaxation modulus *G*(*t*) (i.e., the shear stress relaxation given a unit strain) of a network can be decomposed into a superposition of exponential relaxation modes, each associated with a non-zero eigenvalue of the connectivity matrix (see the detailed derivations in Supplemental Material Section A) [45]:

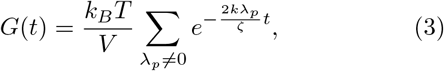

where *V* is the system volume. In this work, all calculations of eigenvalues are based on the connectivity matrix that explicitly includes all *m* beads of each chain. For every pair of chains to be crosslinked, we randomly select one bead from each chain to form the crosslink. From Eq. (3), we compute the zero-shear viscosity as

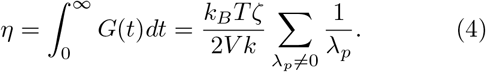

We note that the above viscosity does not include the solvent contribution. We also neglect hydrodynamic interactions [37], which should be a good approximation for solutions with a relatively high polymer volume fraction [46].

To simplify the notation, we set the time unit as *ζ/*2*k*, the viscosity unit as *k*_*B*_*Tζ/*2*V k*, and the stress unit as *k*_*B*_*T/V*. In the following, all variables are dimensionless unless otherwise mentioned. Given the eigenvalues, the storage modulus *G*^′^(*ω*) and loss modulus *G*^′′^(*ω*) can be found via (Supplemental Material Section A)

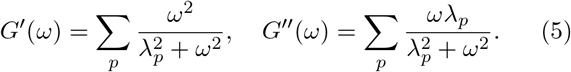

### Eigenvalue distribution for clusters of fixed size

It is clear now that the viscoelastic behaviors of the condensate are encoded in its eigenvalue distribution. Meanwhile, since the condensate consists of many separate clusters, we first seek to understand the eigenvalue distributions for clusters of fixed size.

As an example, let us first consider the formation of a cluster from *n* disconnected chains (Figure 2a). Without any crosslinks, the system’s eigenvalues are identical to those of the original Rouse model (i.e., a linear chain of *m* beads), which consist of *m* distinct modes with eigenvalues,

**FIG. 2.**
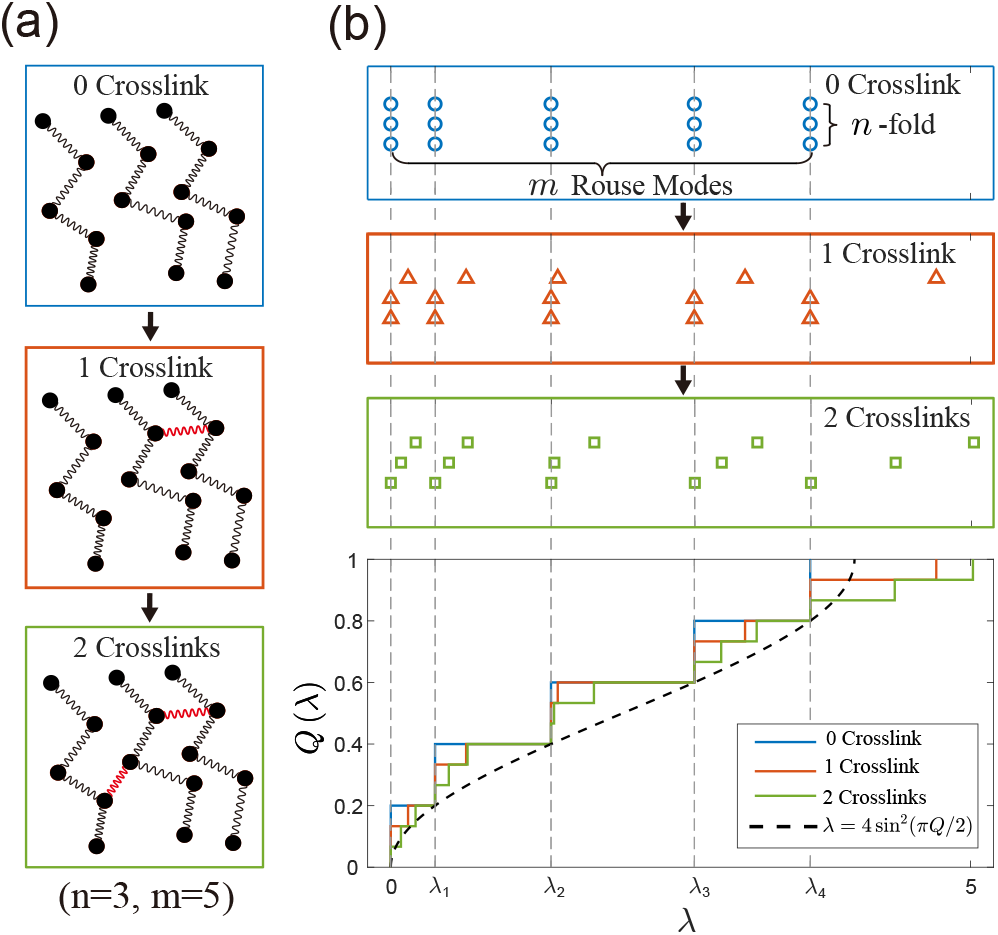
Eigenvalues for a single cluster. (a) A schematic of how a 3-chain cluster is constructed by adding crosslinks one by one. In this example, each chain has *m* = 5 beads. (b) Visualization of the eigenvalues as the number of crosslinks increases, corresponding to (a). The vertical dashed lines mark the *m* distinct Rouse modes (*λ*_0_ through *λ*_4_) of the uncrosslinked chains. The upper panels display the discrete eigenvalues for the initial uncrosslinked chains (blue), a single-crosslink intermediate (red), and the final cluster (green). The bottom panel shows the evolution of the cumulative distribution of the eigenvalues, *Q*(*λ*). The black dashed curve is the analytical relationship *λ* = 4 sin^2^(*πQ/*2) from Eq. (6), which holds exactly for the discrete Rouse modes where *Q*(*λ*_*q*_) = *q/m*.

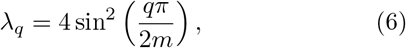

where *q* = 0, 1, …, *m* − 1, which we call the Rouse modes, each with *n*-fold degeneracy (see detailed derivation in Supplemental Material Section C) [35, 38, 47, 48]. As shown in Figure 2a and b, adding a crosslink reduces the degeneracy of each degenerate Rouse mode *λ*_*q*_ by one, while increasing the number of eigenvalues in the interval (*λ*_*q*_, *λ*_*q*+1_) by one according to the interlacing theorem [49] (Supplemental Material Section C). As *n* −1 crosslinks are added to form the final cluster, only one eigenvalue of the original *n*-fold degenerate eigenvalues remains at its initial value *λ*_*q*_. The remaining *n* − 1 are shifted upwards into the interval (*λ*_*q*_, *λ*_*q*+1_) (Figure 2a, b; see Figure S2 for a more general case).

We note that the eigenvalue distribution of a cluster for *λ > λ*_1_ retains the overall shape of a Rouse chain. One can see this by comparing the cumulative eigenvalue distribution *Q*(*λ*) (the fraction of eigenvalues smaller than *λ*) with that of the original Rouse chain as shown in Figure 2b. Intriguingly, for both the uncrosslinked system and the crosslinked cluster, we have *Q*(*λ*_*q*_) = *q/m* as one can see from the eigenvalue distributions in the three top panels in Figure 2b. Indeed, *Q*(*λ*) of the crosslinked clusters coincides with that of the original Rouse chain at *λ*_*q*_ (Figure 2b, bottom panel). Therefore, the cluster’s relaxation behavior below the characteristic Rouse time 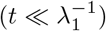 should approximately equal that of a single chain (see a detailed proof in Supplemental Material Section E). On the other hand, it is the *n* − 1 eigenvalues in the interval (0, *λ*_1_), which we call the slow collective modes, that govern the long-term viscoelastic behavior of the cluster and also the entire condensate, as we show in the later section.

We remark that clusters of a given size can exhibit multiple distinct topologies (Figure S5). A key feature of the ER random graph is that the relative likelihood of different topologies for clusters of a given size *n* (i.e., the count of clusters with a specific topology of size *n*, divided by the total count of all clusters of size *n*) is independent of *c*. Therefore, to obtain the eigenvalue distribution for the condensate, one only needs to find the eigenvalue distribution of fixed-size clusters averaged over all possible topologies, which we denote as *p*_*n*_(*λ*) (see numerical details in Supplemental Material Section D3), and then integrate *p*_*n*_(*λ*) over cluster sizes using the cluster size distribution *P* (*n*) (Eq. (1)). Similar to the example in Figure 2, *p*_*n*_(*λ*) has the same overall shape as the original Rouse model for *λ > λ*_1_, as one can see from the cumulative eigenvalue distribution (Figure 3a).

**FIG. 3.**
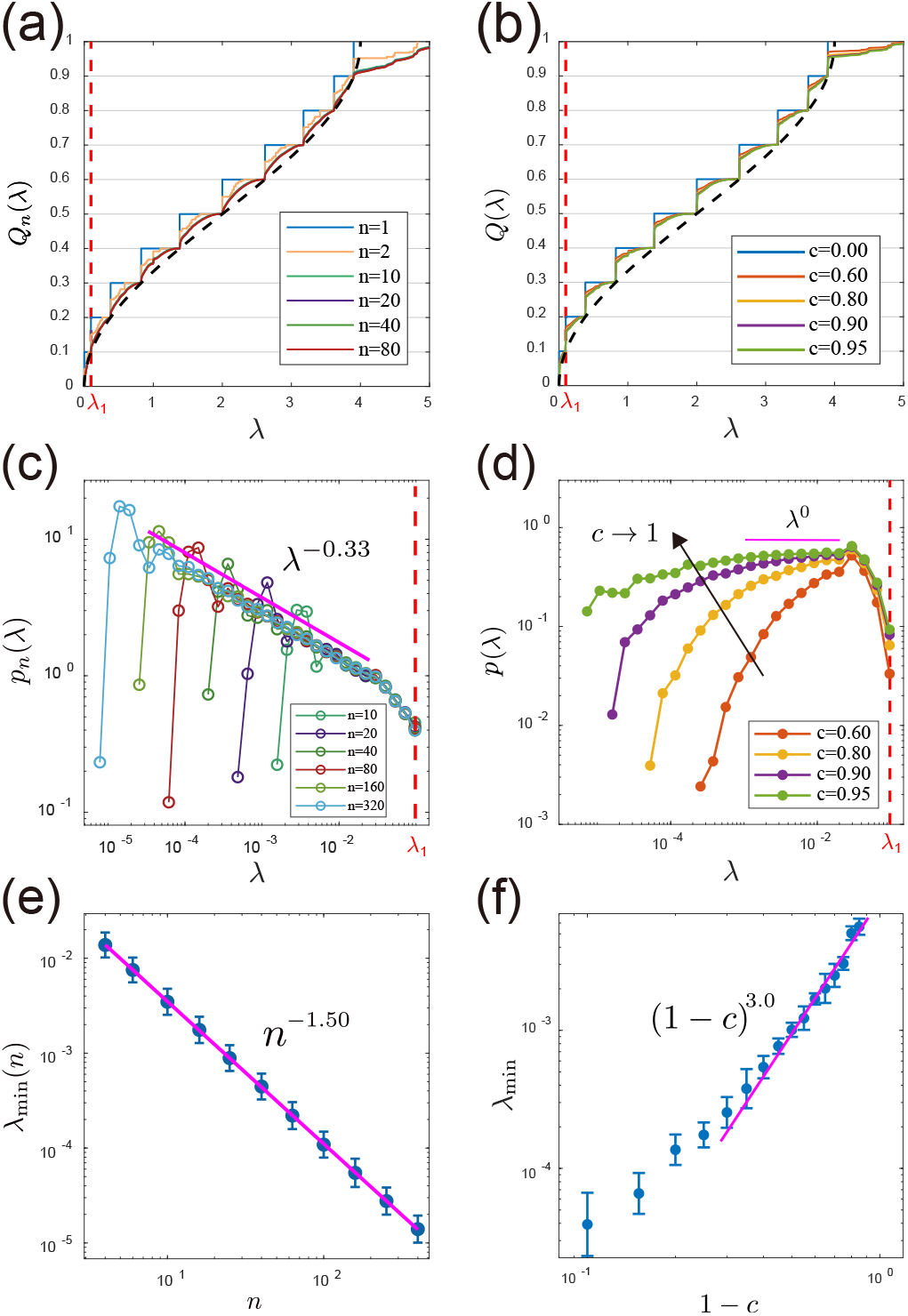
Eigenvalues for clusters of fixed size and the entire condensate. (a) The cumulative eigenvalue distribution functions *Q*(*λ*) for clusters of different sizes *n*. (b) The cumulative eigenvalue distribution functions *Q*(*λ*) for the entire condensate with different mean crosslink numbers per chain, *c*. The case *c* = 0 represents the fully uncrosslinked system composed of multiple Rouse chains. In (a, b), the red dashed line indicates the position of the first Rouse mode *λ*_1_ and the black dashed curve is the analytical relationship *λ* = 4 sin^2^(*πQ/*2) for a single Rouse chain from Eq. (6). (c) The average eigenvalue distribution in the interval (0, *λ*_1_) for clusters of multiple sizes *n*. In (a, c), for each cluster size, we randomly sample 500 labeled topologies. (d) The eigenvalue distribution in the interval (0, *λ*_1_) for the entire condensate, which becomes constant for small *λ* as *c* approaches 1. Here, each curve represents an average over 5 realizations. (e) Scaling of the lower cutoff *λ*_min_ of *p*_*n*_(*λ*) with the cluster size *n*. The error bars show the geometric standard deviation across 100 labeled topologies. (f) Scaling of the lower cutoff *λ*_min_ of the condensate with 1 *c*. For each data point, *λ*_min_ is the geometric mean of the 100 eigenvalues (10 smallest eigenvalues from each of 10 independent realizations), with the error bar indicating the geometric standard deviation. In this figure, *m* = 10 and *N*_*p*_ = 2 × 10^5^ for (b, d, f).

Interestingly, given the self-similar properties of Rouse model, we demonstrate that the eigenvalue distribution must satisfy the scaling form *p*_*n*_(*λ*) = 1*/*(*mλ*_1_)*f*_*n*_(*λ/λ*_1_) within the interval (0, *λ*_1_) (Supplemental Material Section C); more interestingly, we find that *p*_*n*_ exhibits a power-law scaling for small *λ* (Figure 3c):

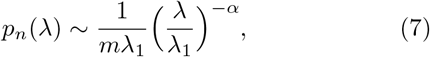

down to a cutoff that scales with the cluster size (Figure 3e),

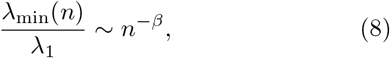

Note that the cluster size *n* is much smaller than the total number of polymer chains *N*_*p*_ due to the upper cutoff *n*^*^ in the cluster size distribution (Eq. (1)).

Given the eigenvalue distribution, one can compute the extra viscosity for clusters of size *n* relative to uncrosslinked systems, which is dominated by the slow modes such that 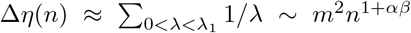 (Supplemental Material Section C). Meanwhile, one can compute the viscosity using its proportionality to the radius of gyration [39], and we find that Δ*η*(*n*) ∼ *m*^2^*n*^3*/*2^ (Supplemental Material Section B). Given the two expressions of the same variable, we obtain an intriguing relation between the exponents:

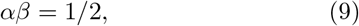

which agrees perfectly with our numerical calculations, where *α* ≈0.33 and *β*≈ 1.50 (Figure 3c, e). In the next section, we show that Eq. (9) implies a constant eigenvalue distribution for small *λ* for the entire condensate.

### From clusters to the entire condensate

We next connect the eigenvalue distribution for clusters of fixed size to that of the entire condensate. We note that the eigenvalue distribution of the entire condensate for eigenvalues that are larger than *λ*_1_ mirrors that of a Rouse chain for the same reason as clusters of fixed size, since the condensate is composed of multiple separate clusters (see Figure 3a, b). However, for the slow modes within the interval (0, *λ*_1_), which set the long-time-scale behavior of the condensate, the eigenvalue distribution is much more interesting.

To obtain the probability distribution between 0 and *λ*_1_ for the entire condensate, we integrate all possible cluster sizes:

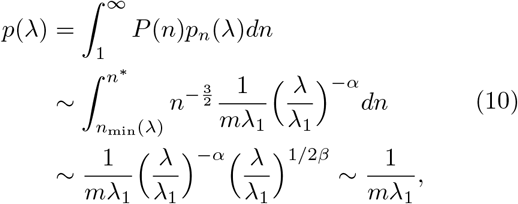

where *n*_min_(*λ*) is the minimum cluster size to observe the target *λ*, which satisfies *n*_min_(*λ*) ∼ (*λ/λ*_1_)^−1*/β*^ according to Eq. (8). *n*^*^ is the upper cutoff in the cluster size distribution (Eq. (1)). Here, we have used the relationship *αβ* = 1*/*2.

Surprisingly, the integration over cluster sizes yields a constant eigenvalue distribution for small *λ* for the entire condensate. Furthermore, the constant distribution has a lower cutoff *λ*_min_, which happens when the lower and upper cutoffs in the integral of Eq. (10) are comparable, i.e., *n*_min_(*λ*_min_) ≈ *n*^*^. This gives:

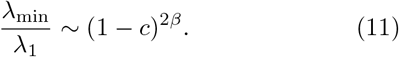

We verify our predictions by numerically simulating the randomly crosslinked condensate. Specifically, we randomly add *N*_*p*_*c/*2 crosslinks to a system of *N*_*p*_ initially disconnected chains (each consisting of *m* beads). We note that since *N*_*p*_ is very large in our simulations, we virtually never see multiple crosslinks between the same two chains. We then compute the eigenvalues of the full connectivity matrix with size *N*_*p*_*m* ×*N*_*p*_*m*. Our theoretical predictions agree perfectly with our numerical calculations (Figures 3d, f), and also with previous calculations based on replica methods for random connectivity matrices [50, 51].

### Viscoelasticity of the condensate

In the following, we compute the relaxation modulus *G*(*t*) using Eq. (3) and the established eigenvalue distribution, accounting for the *N*_*p*_*m* total eigenvalues for the condensate.

For 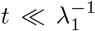, the relaxation modulus of the condensate is governed by eigenvalues larger than *λ*_1_ and resembles that of the original Rouse model due to their similar eigenvalue distributions (Figure 3b; see a detailed proof in Supplemental Material Section E). Considering the limit *m* ≫1, the Rouse model’s eigenvalue distribution becomes *p*(*λ*) ∼ *λ*^−1*/*2^ for *λ*_1_ ≪*λ*≪ 1 (see Eq. (6)) [35, 38]. Therefore, 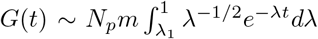. This yields a constant plateau *G*(*t*) ∼ *N*_*p*_*m* for *t* ≪ 1, followed by the characteristic Rouse decay 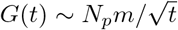 for 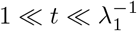 (Figure 4a) [35, 38, 52].

**FIG. 4.**
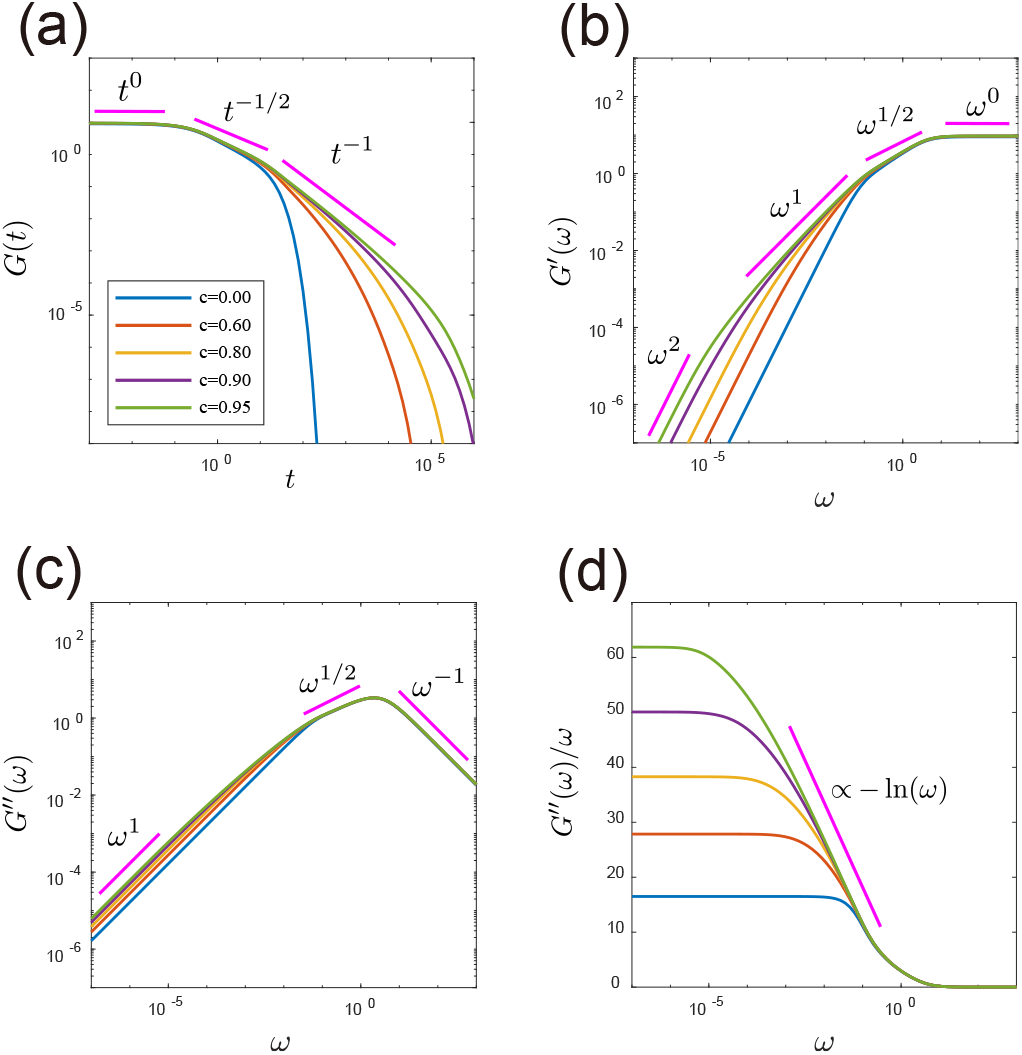
Viscoelasticity of the random crosslinked condensate computed with *N*_*p*_ = 2 ×10^5^ and *m* = 10, averaged over 30 independent realizations. We normalize all moduli in this figure by *N*_*p*_. (a) The relaxation modulus *G*(*t*) as a function of time *t*. (b) The storage modulus *G*^*′*^(*ω*) as a function of angular frequency *ω*. (c) The loss modulus *G*^*′′*^(*ω*) as a function of angular frequency *ω*. (d) *G*^*′′*^(*ω*)*/ω* as a function of angular frequency *ω*.

For 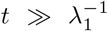, the relaxation is dominated by the constant eigenvalue distribution smaller than *λ*_1_ (Eq. (10)); therefore,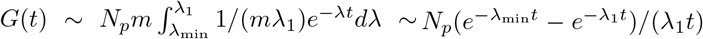. This simplifies to an inverse power-law decay 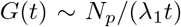 for 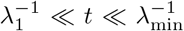, and an exponential decay 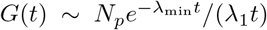 for 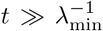. We summarize the temporal scaling of *G*(*t*) for *t* ≫ 1:

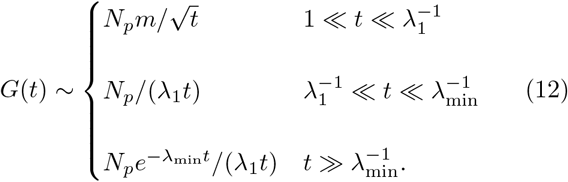

To verify the predicted scaling of the relaxation modulus, we numerically compute the relaxation modulus using the eigenvalues directly computed from the full connectivity matrix of an entire condensate (Figure 3b). The perfect agreement between the predictions and the numerical results strongly supports our theory (Figure 4a). Given the relaxation modulus, one can find the dynamic moduli as 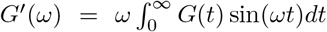 and 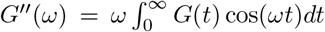 [35, 52]. This yields the following frequency scaling of *G*^′^ and *G*^′′^ as (see detailed derivations in Supplemental Material Section E)

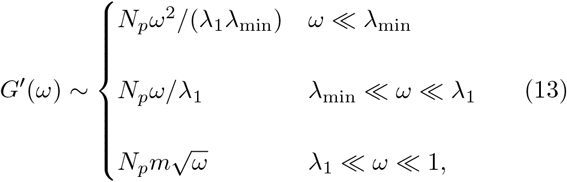

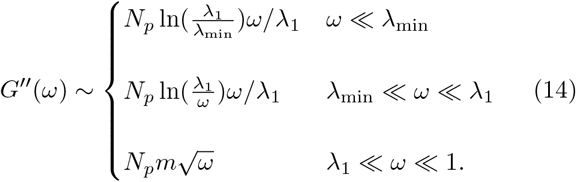

We remark that the anomalous linear frequency scaling of *G*^′^(*ω*) and *G*^′′^(*ω*) (with a logarithmic correction) for *λ*_min_ ≪*ω*≪ *λ*_1_ stems from the constant eigenvalue distribution, reflecting the abundant slow collective modes. The predicted scalings of the dynamic moduli agree well with the computed moduli based on Eq. (5) (Figure 4b, c). From the low-frequency scaling of the loss modulus, we also obtain the scaling of viscosity vs. the distance to gelation threshold (Figure 4d): *η* = lim_*ω*→0_ *G*^′′^(*ω*)*/ω* ∼ − ln(1 −*c*), in agreement with previous works [45]. In the Supplemental Material Section D2, we also include a complete derivation of the viscosity over the full range 0 *< c <* 1.

Notably, the enhancement of storage modulus due to crosslinks at low frequency *ω < λ*_1_ is much more significant than that of the loss modulus (comparing Figure 4b and c). To understand the boost of storage modulus, one can consider the low frequency limit *ω* ≪ *λ*_min_ where 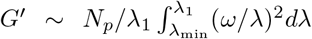, while 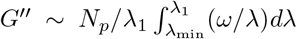 (see Eq. (5)). Due to its stronger *λ*^−2^ dependence, the storage modulus is more responsive to the slow modes.

### Comparison with experimental data

We validate our predictions by comparing the predicted dynamic moduli with experimental data obtained from diverse protein condensates [11, 12, 34]. We rescale the experimental data by the crossover point (*ω*_0_, *G*_0_), where *G*^′^(*ω*_0_) ≈ *G*^′′^(*ω*_0_) ≈*G*_0_. We then align the theoretical curve with this normalized data, shifting it on the log-log plot to achieve the best overall agreement. Our theoretical curves with parameters *m* = 5 and *c* = 0.95 align reasonably well with all the experimental data, even though the underlying components of these condensates are entirely different (Figure 5). We note that the results are not sensitive to the exact value of *c* as long as it is close to 1, and the bead number *m* primarily alters the high-frequency fit, while the low-frequency regime (*ω/ω*_0_ *<* 1) remains robust for all *m* (see Supplemental Material Section F). Our model successfully captures the linear scaling of *G*^′^ and *G*^′′^ at low frequencies. The universality of this anomalous scaling across diverse condensates suggests that the soft modes involving multiple chains in a random crosslinked polymer solution are general mechanisms of stress relaxation.

**FIG. 5.**
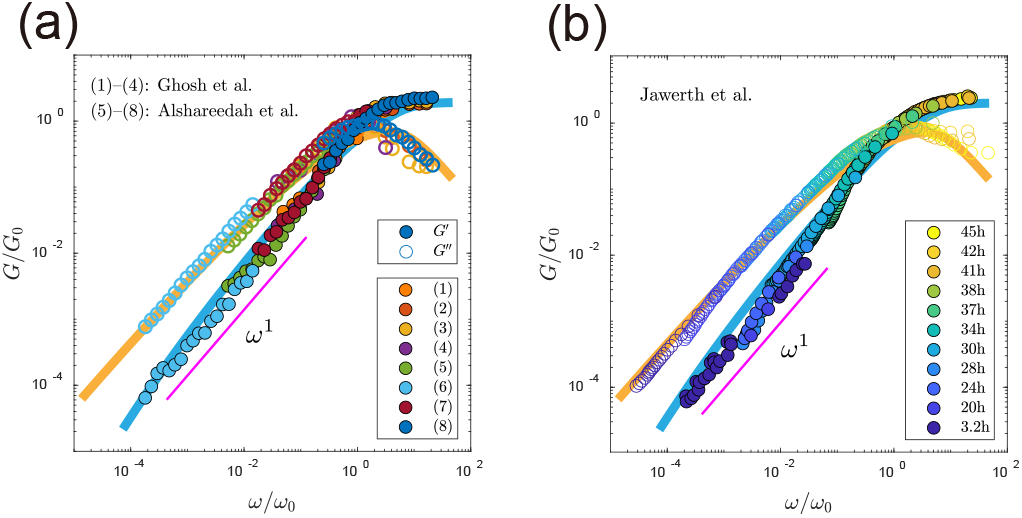
Comparison between theoretical predictions and experimental data. (a) Data from multiple studies: (1)-(4) condensates formed by binary mixture [12]; (5)-(8) condensates formed by different variants of the intrinsically disordered prion-like low-complexity domain of the RNA-binding protein hnRNPA1 [34]. (b) Condensates formed by protein PGL-3 at different waiting times after sample formation [11]. The solid lines are the theoretical predictions (blue for *G*^*′*^, yellow for *G*^*′′*^) with fixed parameters *m* = 5 and *c* = 0.95. The experimental data are normalized by the crossover point where *G*^*′*^(*ω*_0_) ≈*G*^*′′*^(*ω*_0_) ≈ *G*_0_. The magenta line has a slope of 1 in the log-log plot.

## Discussion

In this work, we study a minimal model of biomolecular condensates as crosslinked polymer solutions below gelation. The resulting network topology is captured by a sub-critical Erdős-Rényi random graph, which decomposes the system into an ensemble of clusters of varying sizes *n*. Our analysis of clusters of size *n* reveals that the eigenvalue distribution exhibits a power-law scaling for slow modes between 0 and *λ*_1_, *p*_*n*_(*λ*) ∼*λ*^−*α*^ down to a size-dependent cutoff *λ*_min_ ∼*n*^−*β*^. We further derive a universal relation *αβ* = 1*/*2, in agreement with our simulations where *α* ≈ 0.33, *β* ≈ 1.50.

Going from single clusters to the entire condensate, we prove that the eigenvalue distribution of the whole condensate must be constant with a lower cutoff *λ*_min_ ∼ (1 −*c*)^2*β*^. The abundance of slow modes leads to the anomalous scaling of the storage and loss moduli: *G*^′^(*ω*) ∼ *ω* and *G*^′′^(*ω*) ∼ *ω* ln(*λ*_1_*/ω*) in the frequency regime *λ*_min_ ≪*ω* ≪*λ*_1_. Notably, the storage modulus is significantly higher in the random crosslinked system than in the uncrosslinked system at low frequencies.

Our model is simplified, and real-world condensates can be more complex due to factors such as specific molecular interactions. Yet, our predictions from this minimal model are in remarkable agreement with experimental data from diverse biomolecular condensates, including binary-mixture condensates, sequence-variant condensates, and aging protein condensates (Figure 5). This indicates that the slow collective modes generated by random connectivity provide a universal mechanism for the anomalous linear scaling of the storage modulus at low frequencies. We note that across all the experimental data we analyze, the storage modulus is lower than the loss modulus at low frequencies, indicating that the condensates are liquid over long timescales, which is why we focus primarily on condensates below gelation. Exploring how deviations from our minimal model, such as effects of loops, gelation, and network structures [53], would change the viscoelastic response is an important direction for future research. Detailed analytical derivations, numerical procedures, and full methodologies are provided in the Supplementary Information (SI) Appendix.

We thank Huan-Xiang Zhou for discussions related to this work. The research was funded by the National Natural Science Foundation of China (Grant No. 12474190), the National Key Research and Development Program of China (2024YFA0919600), and grants from the Peking-Tsinghua Center for Life Sciences.

