## Supplemental Material for "Random crosslinks generate anomalous scaling of dynamic moduli of biomolecular condensates"

(Dated: June 2, 2026)

### CONTENTS

A. Dynamic Modulus of a Generalized Rouse Network 2
B. Viscosity for a Cluster of size $n$ 3
1. Relation Between Viscosity and Radius of Gyration 3
2. Relation Between the Viscosity of a Cluster and the Viscosity of its Coarse-grained Graph 4
3. Calculation of the Viscosity for Clusters of Fixed Size 5
C. Eigenvalue Distribution for Clusters of Size $n$ 6
D. Properties of an ER Random Graph 9
1. Physical Justification and Derivation of the Branch Size Distribution 9
2. Viscosity of an ER Random Graph 10
3. Cluster Topology Distribution of Given Size and Its Sampling Procedure 12
E. Derivation of Dynamic Moduli 12
1. High-Frequency Limit ($\omega \gg 1$) 14
2. Rouse Regime ($\lambda_1 \ll \omega \ll 1$) 14
3. Collective-Behavior Regime ($\lambda_{\min} \ll \omega \ll \lambda_1$) 15
4. Low-Frequency Limit ($\omega \ll \lambda_{\min}$) 15
F. Parameter Selection and Experimental Fitting 16

References 16

#### A. Dynamic Modulus of a Generalized Rouse Network

According to the generalized Rouse model for a connected network of $N$ beads with arbitrary configurations [1–3], the position of bead $i$ follows the Langevin equation:

$$\zeta \dot{\boldsymbol{R}}_i = -kL_{ij}\boldsymbol{R}_j + \boldsymbol{\xi}_i \tag{S1}$$

Here, $\zeta$ is the friction coefficient and $k$ is the spring constant, which can be written as $k = 3k_BT/b^2$ where $k_B$ is the Boltzmann constant, $T$ is the temperature, and $b^2$ is the mean square distance between two connected beads. The random force $\boldsymbol{\xi}_i$ is generated by thermal fluctuation and satisfies the fluctuation-dissipation theorem (FDT) [4],

$$\langle \xi_{i,\alpha}(t)\xi_{j,\beta}(t')\rangle = 2\zeta k_BT\delta_{ij}\delta_{\alpha\beta}\delta(t-t'). \tag{S2}$$

Here $\delta_{ij}$ and $\delta_{\alpha\beta}$ are Kronecker delta functions, and $\delta(t-t')$ is Dirac delta function. The Greek letters represent the directions in the Cartesian coordinate system. The connectivity matrix $L_{ij}$ is a $N \times N$ symmetric matrix defined as $L_{ij} = \delta_{ij}\sum_k A_{ik} - A_{ij}$, where $A_{ij}$ is the adjacency matrix with $A_{ij} = 1$ if beads $i$ and $j$ are connected and 0 otherwise. The connectivity matrix describes the connectivity between beads with non-negative eigenvalues $\lambda_p \geq 0$ and orthonormal real eigenvectors $|u_p\rangle$. We project the $N \times 3$ position vector $|\boldsymbol{R}\rangle$ onto the eigenmodes such that $\boldsymbol{c}_p = \langle u_p|\boldsymbol{R}\rangle$, a $1 \times 3$ vector that satisfies

$$\zeta \dot{\boldsymbol{c}}_p = -k\lambda_p \boldsymbol{c}_p + \boldsymbol{f}_p, \tag{S3}$$

where $\boldsymbol{f}_p = \langle u_p|\boldsymbol{\xi}\rangle$, which also satisfies the FDT,

$$\langle f_{p,\alpha}(t)f_{q,\beta}(t')\rangle = 2\zeta k_BT\delta_{pq}\delta_{\alpha\beta}\delta(t-t'). \tag{S4}$$

We introduce a step shear with strain $\gamma$ at $t = 0$. The shear deformation is applied along the x-direction with a gradient along the y-direction, transforming the bead positions as:

$$x_i(0^+) = x_i(0^-) + \gamma y_i(0^-), \quad y_i(0^+) = y_i(0^-), \quad z_i(0^+) = z_i(0^-). \tag{S5}$$

The shear stress $\sigma_{xy}$, defined as force per unit area, can be written as [5]

$$\sigma_{xy} = -\frac{1}{V}\sum_i F_{i,x}y_i = \frac{k}{V}\langle x|L|y\rangle, \tag{S6}$$

where $F_{i,x}$ is the total force on bead $i$ in the x-direction and $V$ is the system volume. Decomposing the position vectors using the eigenvectors of $L$, where $|x\rangle = \sum_p c_{px}|u_p\rangle$ and $|y\rangle = \sum_q c_{qy}|u_q\rangle$, we obtain

$$\sigma_{xy} = \frac{k}{V}\sum_{p,q} c_{px}c_{qy}\langle u_p|L|u_q\rangle = \frac{k}{V}\sum_p \lambda_p c_{px}c_{py}, \tag{S7}$$

where we use the orthogonality $\langle u_p|u_q\rangle = \delta_{pq}$ and the eigenvalue equation $L|u_p\rangle = \lambda_p|u_p\rangle$. Under the step strain at $t = 0$, the initial conditions can be rewritten as

$$\begin{aligned} c_{py}(0^+) &= c_{py}(0^-), \\ c_{px}(0^+) &= c_{px}(0^-) + \gamma c_{py}(0^-). \end{aligned} \tag{S8}$$

This leads to the initial cross-correlation:

$$\langle c_{px}(0^+)c_{py}(0^+)\rangle = \gamma\langle c_{py}^2\rangle = \gamma\frac{k_BT}{k\lambda_p}, \tag{S9}$$

where the equilibrium variance $\langle c_{py}^2\rangle = k_BT/(k\lambda_p)$ follows from equipartition theorem. Taking the time derivative of $\langle c_{px}c_{py}\rangle$ and using Eq. (S3):

$$\frac{d}{dt}\langle c_{px}c_{py}\rangle = \langle \dot{c}_{px}c_{py}\rangle + \langle c_{px}\dot{c}_{py}\rangle = -\frac{2k\lambda_p}{\zeta}\langle c_{px}c_{py}\rangle, \tag{S10}$$

leading to

$$\langle c_{px}(t)c_{py}(t)\rangle = \gamma\frac{k_BT}{k\lambda_p}e^{-(2k\lambda_p/\zeta)t}, \tag{S11}$$

and the time-dependent shear stress

$$\langle\sigma_{xy}(t)\rangle = \frac{k}{V}\sum_{p,\,\lambda_p\neq 0}\lambda_p\left(\gamma\frac{k_BT}{k\lambda_p}e^{-(2k\lambda_p/\zeta)t}\right) = \frac{\gamma k_BT}{V}\sum_{p,\,\lambda_p\neq 0}e^{-t/\tau_p}, \tag{S12}$$

where $\tau_p \equiv \zeta/(2k\lambda_p)$ is the relaxation time of mode $p$. The shear relaxation modulus $G(t)$ is then:

$$G(t) \equiv \frac{\langle\sigma_{xy}(t)\rangle}{\gamma} = \frac{k_BT}{V}\sum_{p,\,\lambda_p\neq 0}e^{-t/\tau_p}. \tag{S13}$$

The frequency-dependent complex shear modulus $G^*(\omega) = G'(\omega) + iG''(\omega)$ is obtained from the Fourier transform of the relaxation modulus:

$$G^*(\omega) = i\omega\int_0^\infty G(t)e^{-i\omega t}dt. \tag{S14}$$

Substituting Eq. (S13) into the transform yields:

$$G^*(\omega) = i\omega\frac{k_BT}{V}\sum_{p,\,\lambda_p\neq 0}\int_0^\infty e^{-t/\tau_p}e^{-i\omega t}dt = \frac{k_BT}{V}\sum_{p,\,\lambda_p\neq 0}\frac{i\omega\tau_p}{1+i\omega\tau_p}. \tag{S15}$$

Separating the real and imaginary parts of Eq. (S15) gives the storage modulus $G'(\omega)$ and the loss modulus $G''(\omega)$:

$$\begin{aligned} G'(\omega) &= \frac{k_BT}{V}\sum_{p,\,\lambda_p\neq 0}\frac{\omega^2\tau_p^2}{1+\omega^2\tau_p^2}, \\ G''(\omega) &= \frac{k_BT}{V}\sum_{p,\,\lambda_p\neq 0}\frac{\omega\tau_p}{1+\omega^2\tau_p^2}. \end{aligned} \tag{S16}$$

### B. Viscosity for a Cluster of size $n$

#### *1. Relation Between Viscosity and Radius of Gyration*

The viscosity $\eta$ is defined as the time integral of relaxation modulus after a step strain: $\eta = \int_0^\infty \langle\sigma_{xy}(t)\rangle dt/\gamma$. Using Eq. (S12), the viscosity can be expressed as the sum of the inverse of all nonzero eigenvalues:

$$\eta = \frac{\zeta k_BT}{2kV}\sum_p\frac{1}{\lambda_p}. \tag{S17}$$

The mean-square radius of gyration quantifies the spatial size of a macromolecule, which can be computed as $\langle R_g^2\rangle = \sum_{i,j}\langle(\boldsymbol{R}_i - \boldsymbol{R}_j)^2\rangle/2N^2$ [6], where $N$ is the number of beads. Here, the average is over all possible configurations of the molecule at thermal equilibrium. We rewrite $\boldsymbol{R}_i - \boldsymbol{R}_j = \sum_p \boldsymbol{c}_p(u_{pi} - u_{pj})$, and it is convenient to consider the fluctuation in one direction:

$$\begin{aligned} \sum_{i,j}\langle(x_i - x_j)^2\rangle &= \langle\sum_{p,q}c_{px}c_{qx}\sum_{i,j}(u_{pi}-u_{pj})(u_{qi}-u_{qj})\rangle \\ &= \sum_p\langle c_{px}^2\rangle\sum_{i,j}(u_{pi}^2 + u_{pj}^2 - 2u_{pi}u_{pj}) \\ &= \sum_p\frac{2k_BT}{k\lambda_p}[N\sum_i u_{pi}^2 - (\sum_i u_{pi})^2] \\ &= \sum_p\frac{2Nk_BT}{k\lambda_p}. \end{aligned} \tag{S18}$$

In the above derivation, we use the fact that the fluctuation of each eigenmode is independent, as well as the equipartition theorem for $\langle c_{px}^2 \rangle$. We also use the normalization condition: $\sum_i u_{pi}^2 = 1$, and the condition, $\sum_i u_{pi} = 0$ since rigid-body motion is not included. Since the three directions are equivalent, we obtain

$$\langle R_g^2 \rangle = \frac{3k_BT}{Nk} \sum_p \frac{1}{\lambda_p} = \frac{b^2}{N} \sum_p \frac{1}{\lambda_p}. \tag{S19}$$

Combining (S17) and (S19), we recover the established relation [2]:

$$\eta = \frac{N\zeta}{6V} \langle R_g^2 \rangle. \tag{S20}$$

#### *2. Relation Between the Viscosity of a Cluster and the Viscosity of its Coarse-grained Graph*

In the remainder of this section and also the rest of the Supplemental Material, we use dimensionless variables to simplify the notation with the same units we introduce in the main text. In particular, converting the viscosity and mean-square radius of gyration to the dimensionless forms we use in the main text, we find

$$\eta = N\langle R_g^2 \rangle = \sum_p \frac{1}{\lambda_p}, \tag{S21}$$

where the viscosity unit is $k_BT\zeta/(2Vk)$ and the length unit is $b$.

The advantage of Eq. (S21) is that one can convert the problem of finding viscosity (i.e., finding all the eigenvalues of the connectivity matrix $L$) to the problem of calculating the mean-square radius of gyration. In particular, for clusters, the Kramers theorem significantly simplifies the calculation of the radius of gyration: $\langle R_g^2 \rangle = \sum_e \pi_e / N^2$, where $\pi_e$ is the number of all strands between different beads that pass through a given edge [6]. Therefore, one can compute the viscosity as

$$\eta = \sum_e \pi_e / N. \tag{S22}$$

In the following, we apply the relationship between viscosity and radius of gyration to a cluster. Consider a cluster of $n$ linear chains with $n-1$ crosslinks, each containing $m$ beads such that the total number of beads $N = nm$. For every crosslinked pair of chains, we randomly select one bead from each chain to add the crosslink. The network topology is defined by the coarse-grained graph $G$, where each node represents a linear chain and edges represent crosslinks between chains. We compute the viscosity $\eta$ of the complete network and separate the edges into two categories $nm\eta = \sum_{e_c} \pi_{e_c} + \sum_{e_b} \pi_{e_b}$ where $e_c$ are crosslinks between polymer chains and $e_b$ denotes backbone bonds within each linear chain. We calculate these two terms separately as follows.

*Crosslink contribution*—For crosslinks, each $e_c$ corresponds to a coarse-grained edge $e_c'$ in graph $G$ with $\pi_{e_c} = m^2 \pi_{e_c'}^G$ (see Figure 1b in the main text), leading to the total contribution $\sum_{e_c} \pi_{e_c} = m^2 n \eta_G$ where $\eta_G$ is the viscosity of the coarse-grained graph $G$.

*Backbone contribution*—For each chain $s$, the backbone contribution $\sum_{e_b \in s} \pi_{e_b}$ is equal to the sum, over all pairs of beads $(i,j)$, of the length of the path segment between $i$ and $j$ that lies within chain $s$ (see the yellow paths in Figure S1). This total path length within $s$ can be decomposed into three categories based on the locations of the endpoints $i$ and $j$ (Figure S1): (1) paths with both endpoints in $s$, (2) paths with one endpoint in $s$ and the other outside $s$, (3) paths with both endpoints outside $s$.

In the following, we compute the three categories of backbone contribution separately:

(1) The sum of path distances over all pairs with both endpoints in $s$ is simply $\frac{1}{2}\sum_{i=1}^m \sum_{j=1}^m |i-j| = \sum_{i=0}^{i=m} i(m-i) = m\eta_p$ (Figure S1a), where $\eta_p = (m^2-1)/6$ is the viscosity of a single linear chain with $m$ beads. Summing over all $n$ chains gives the contribution $nm\eta_p$.

(2) The second term represents the sum of path lengths traversing bonds in $s$ for all pairs with one endpoint in $s$ and the other outside ($i \in s, j \notin s$) (Figure S1b). The total number of such bead pairs is $m \times (n-1)m$. For each pair $(i,j)$, the average path distance on chain $s$ (connecting $j$ to the branch attachment point $j'$) is given by: $\bar{d} = \frac{1}{m^2}\sum_{i=1}^m \sum_{j'=1}^m |i-j'| = 2\eta_p/m$. Summing up all $n$ chains gives the contribution $m^2(n-1) \times \bar{d} \times n = 2nm(n-1)\eta_p$.

(3) To calculate the third term, we introduce a branch size function $[s,l]$ as follows: for two linear chains $s$ and $l$ connected by a crosslink, $[s,l]$ is the number of chains of the branch containing $s$ when the crosslink is removed. By

definition, the branch size function satisfies $[s,l]+[l,s]=n$, and $\sum_l [l,s]=n-1$. For each pair of chains $l_1 \neq l_2$ directly connected to $s$, the number of path pairs with end points $(i,j)$ ($i$, $j$ do not have to be in chain $l_1$ or $l_2$) is determined by the size of their respective branches (Figure S1c). There are $m[l_1,s]$ beads in branch $l_1$ and $m[l_2,s]$ beads in branch $l_2$, yielding $m^2[l_1,s][l_2,s]$ possible bead pairs. Thus, the expectation of the sum of path lengths traversing bonds in $s$ for the third term is $\bar{d}\sum_{l_1<l_2} m^2[l_1,s][l_2,s] = 2\eta_p m \sum_{l_1<l_2}[l_1,s][l_2,s]$.

We compute the double summation using the branch size properties and definition of viscosity:

$$\begin{aligned}
\sum_s \sum_{l_1<l_2}[l_1,s][l_2,s] &= \frac{1}{2}\sum_s\sum_{l_1}[l_1,s]\left(\sum_{l_2\neq l_1}[l_2,s]\right) \\
&= \frac{1}{2}\sum_s\sum_{l_1}[l_1,s]\left(\sum_{l_2}[l_2,s]-[l_1,s]\right) \\
&= \frac{1}{2}\sum_s\sum_{l_1}[l_1,s]\left((n-1)-(n-[s,l_1])\right) \\
&= \frac{1}{2}\sum_s\sum_{l_1}[l_1,s]\left([s,l_1]-1\right) \\
&= \frac{1}{2}\sum_s\sum_{l_1}[l_1,s][s,l_1]-\frac{1}{2}\sum_s(n-1) \\
&= \frac{1}{2}\sum_s\sum_{l_1}[l_1,s][s,l_1]-\frac{n(n-1)}{2} \\
&= \eta_G n-\frac{n(n-1)}{2}.
\end{aligned} \tag{S23}$$

In the final step, we use the formula Eq. (S22) to compute the viscosity for the coarse-grained graph $G$. Hence the total contribution for the third term is: $2\eta_p m[\eta_G n - n(n-1)/2]$. Combining crosslink contribution and the three terms in backbone contribution, we obtain the exact formula for the viscosity of a cluster:

$$\eta = n\eta_p + \frac{m^2+3m-1}{3}\eta_G. \tag{S24}$$

In the above equation, the first term on the right side represents the viscosity of $n$ disconnected chains, and the second term represents the extra viscosity generated by crosslinks. The above formula shows that to calculate the viscosity of a cluster of $n$ chains (each with $m$ beads), one only needs to compute the viscosity of the coarse-grained network, which we calculate in the following for clusters of fixed size.

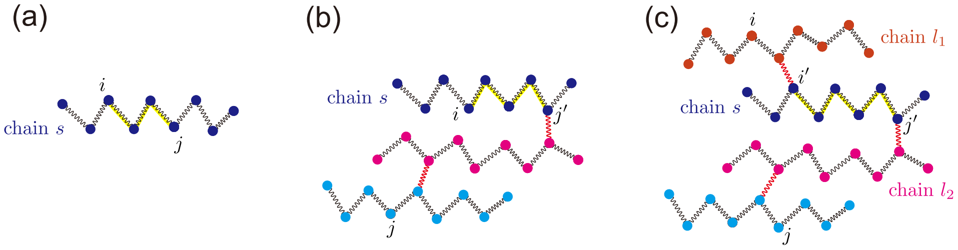


FIG. S1. Illustration of the three categories of paths due to backbone contribution. (a) Case 1: paths with both endpoints $(i,j)$ in chain $s$. (b) Case 2: paths with one endpoint ($i$) in chain $s$ and the other ($j$) outside; $j'$ is the connection point in $s$. (c) Case 3: paths with both endpoints $(i,j)$ outside chain $s$. Their path traverses $s$, entering at point $i'$ and exiting at point $j'$. In all cases, the yellow-highlighted segment on chain $s$ (dark blue) represents the portion of the path that lies within $s$ and contributes to the backbone viscosity.

#### *3. Calculation of the Viscosity for Clusters of Fixed Size*

In this section, we calculate the average viscosity of clusters formed by $n$ chains and $n-1$ crosslinks. As established in the previous section, the viscosity increment $\Delta\eta(n) = \eta - n\eta_p$ relative to an uncrosslinked system is proportional to the viscosity of the coarse-grained graph, $\eta_G(n)$ (Eq. (S24)).

Therefore, we first compute the average viscosity $\langle \eta_G(n) \rangle$ for an ensemble of clusters of size $n$ with different topologies (see the topological sampling procedure in Supplemental Material Section D3). This coarse-grained viscosity corresponds to a network where each polymer chain is treated as a single node. We consider a labeled cluster with $n$ nodes. For any edge connecting nodes $a$ and $b$, we divide the cluster into two components, each with $i$ and $j$ nodes, respectively ($i + j = n - 2$), excluding nodes $a$ and $b$. We then use Eq. (S22) and get the number of paths that pass through the edge, $\pi_e = (i + 1)(j + 1)$. Note that the number of ways to choose $i$ nodes from the $n - 2$ nodes is $(i + j)!/(i!j!)$. For each such partition, the number of distinct labeled clusters with this specific edge division is $(i+1)^{i-1}(j+1)^{j-1}$, using the fact that the number of labeled clusters of size $n$ is $n^{n-2}$ by Cayley's formula [7], where all nodes are distinguishable. Summing over all possible partitions and all $n(n-1)/2$ possible edges, the total sum of $\pi_e$ values for all clusters of size $n$ is:

$$\sum_{\text{all clusters}} \sum_e \pi_e = \frac{n(n-1)}{2} \sum_{i+j=n-2} \frac{(i+j)!}{i!j!}(i+1)^i(j+1)^j = \frac{n!}{2} \sum_{i+j=n-2} \frac{(i+1)^i(j+1)^j}{i!j!}. \tag{S25}$$

Since there are $n^{n-2}$ distinct labeled clusters, the average viscosity becomes:

$$\langle \eta_G(n) \rangle = \frac{1}{n^{n-2}} \cdot \frac{1}{n} \sum_{\text{all clusters}} \sum_e \pi_e = \frac{n!}{2n^{n-1}} \sum_{i+j=n-2} \frac{(i+1)^{(i+1)}(j+1)^{(j+1)}}{(i+1)!(j+1)!}. \tag{S26}$$

Applying Stirling's approximation $m! \approx \sqrt{2\pi m}(m/e)^m$ leads to

$$\frac{(i+1)^{(i+1)}(j+1)^{(j+1)}}{(i+1)!(j+1)!} \approx \frac{e^n}{2\pi\sqrt{(i+1)(j+1)}}. \tag{S27}$$

The sum then becomes

$$\sum_{i+j=n-2} \frac{e^n}{2\pi\sqrt{(i+1)(j+1)}} \approx \frac{e^n}{2\pi} \sum_{k=1}^{n-1} \frac{1}{\sqrt{k(n-k)}} \approx \frac{e^n}{2}. \tag{S28}$$

Substituting back and applying Stirling's approximation to $n!$, we get the dimensionless viscosity:

$$\langle \eta_G(n) \rangle \approx \frac{n!}{2n^{n-1}} \cdot \frac{e^n}{2} \approx \frac{\sqrt{2\pi n}(n/e)^n}{2n^{n-1}} \cdot \frac{e^n}{2} = \sqrt{\frac{\pi}{8}} n^{3/2}. \tag{S29}$$

The coarse-grained viscosity can now be used to determine the viscosity increment for the cluster. By substituting the coarse-grained graph viscosity from Eq. (S29) into the relation from Eq. (S24), we obtain the average viscosity increment for a cluster of size $n$ formed by precursor chains of length $m$:

$$\langle \Delta\eta(n) \rangle \approx \frac{m^2 + 3m - 1}{3} \sqrt{\frac{\pi}{8}} n^{3/2} \sim m^2 n^{3/2}. \tag{S30}$$

#### C. Eigenvalue Distribution for Clusters of Size $n$

To elucidate the eigenvalue distribution for clusters of fixed size $n$, we introduce the eigenvalue interlacing theorem [8]. This theorem describes how the eigenvalues of a graph's Laplacian matrix (i.e., connectivity matrix) change upon the addition of an extra edge (i.e., an extra crosslink). If we denote the Laplacian eigenvalues sorted in non-decreasing order before adding the crosslink as $\{\alpha_i\}$ and after as $\{\gamma_i\}$, the theorem states that they will interlace as follows:

$$\alpha_1 \le \gamma_1 \le \alpha_2 \le \gamma_2 \le \cdots \le \alpha_N \le \gamma_N. \tag{S31}$$

In other words, the i-th eigenvalue of the new graph, $\gamma_i$, is located in the interval between the i-th and (i+1)-th eigenvalues of the original graph, $[\alpha_i, \alpha_{i+1}]$. Numerically, we virtually always find $\alpha_i < \gamma_i < \alpha_{i+1}$ if $\alpha_i \neq \alpha_{i+1}$ in this work.

In the following, we consider the formation of a cluster from $n$ disconnected chains, each with $m$ beads (Figure S2 and Figure 2 in the main text). Initially, the system of $n$ disconnected chains is described by a connectivity matrix (or graph Laplacian), $\mathbf{L}_{\text{initial}}$, which has a block-diagonal structure:

$$\mathbf{L}_{\text{initial}} = \begin{pmatrix} \mathbf{L}_{\text{chain}} & \mathbf{0} & \cdots & \mathbf{0} \\ \mathbf{0} & \mathbf{L}_{\text{chain}} & \ddots & \vdots \\ \vdots & \ddots & \ddots & \mathbf{0} \\ \mathbf{0} & \cdots & \mathbf{0} & \mathbf{L}_{\text{chain}} \end{pmatrix}. \tag{S32}$$

Here, each of the $n$ blocks on the diagonal, $\mathbf{L}_{\text{chain}}$, is the $m \times m$ connectivity matrix for a single Rouse chain (i.e., a path graph), which is given by:

$$\mathbf{L}_{\text{chain}} = \begin{pmatrix} 1 & -1 & 0 & \cdots & 0 \\ -1 & 2 & -1 & \cdots & 0 \\ 0 & -1 & 2 & \ddots & \vdots \\ \vdots & \vdots & \ddots & \ddots & -1 \\ 0 & 0 & \cdots & -1 & 1 \end{pmatrix}. \tag{S33}$$

The block-diagonal form of $\mathbf{L}_{\text{initial}}$ directly implies that the uncrosslinked system has the same eigenvalues as those of $\mathbf{L}_{\text{chain}}$. This gives rise to $m$ distinct modes, known as the $q$-th Rouse modes, each with an $n$-fold degeneracy [6, 9, 10]:

$$\lambda_q = 4\sin^2\left(\frac{q\pi}{2m}\right), \quad \text{where } q = 0, 1, \ldots, m-1. \tag{S34}$$

These $\lambda_q$ values are precisely the eigenvalues of the single-chain connectivity matrix $\mathbf{L}_{\text{chain}}$. As $n-1$ crosslinks are added one by one to form the cluster, the interlacing theorem is applied at each step. The final result of this iterative process is that the original $n$-fold degeneracy of each mode $\lambda_q$ is resolved: one eigenvalue remains at its initial position $\lambda_q$, while the remaining $n-1$ are shifted upwards into the interval $(\lambda_q, \lambda_{q+1})$. For the mode $q = m-1$, these $n-1$ eigenvalues are shifted upwards into the interval $(\lambda_{m-1}, \infty)$. As we show in the main text, $Q(\lambda_q) = q/m$ is true for both the uncrosslinked system and the crosslinked cluster (Figure S2); therefore, the cluster's relaxation behavior below the characteristic Rouse time $(1/\lambda_1)$ is essentially indistinguishable from that of a single chain. This conclusion is justified through a rigorous error analysis in Section E.

In the following, we demonstrate that the eigenvalue distribution for clusters of fixed size must satisfy the scaling form, $p_n(\lambda) = f(\lambda/\lambda_1)/(m\lambda_1)$ for $0 < \lambda < \lambda_1$. To show this, let us consider the dynamical equation of the Rouse model with physical units (Eq. (S1)). We remark that coarse-graining the number of beads in a linear chain, $m$, should not change the physics in the collective behavior regime $(0, \lambda_1)$ when $m$ is sufficiently large. As $m$ changes, the friction coefficient and the spring constant change accordingly and scale as $\zeta \sim m^{-1}$ and $k \sim m$: coarse-graining several beads into one bead makes the effective spring constant smaller and the friction coefficient felt by one bead larger [6]. Meanwhile, the relaxation modulus in physical units beyond the Rouse time, $G(t) \approx (k_B T/V)\sum_{i=1}^{n-1}\exp(-2k\mu_i t/\zeta)$ where $\mu_i$ are the $n-1$ eigenvalues in $(0, \lambda_1)$, must remain invariant under changes in the coarse-graining level of $m$, leading to the relation $\mu_i \sim \zeta/k \sim m^{-2} \sim \lambda_1$. Consequently, the normalized eigenvalues $\mu_i/\lambda_1$ must be independent of $m$. We confirm this scaling numerically for a fixed $n = 20$: plotting the rescaled quantities $mQ_n(\lambda)$ versus $\lambda/\lambda_1$ collapses all curves for $m \gtrsim 10$ onto a single master curve (see Figure S3).

The above analysis suggests the following scaling form for the probability distribution of eigenvalues between 0 and $\lambda_1$, $mp_n(\lambda) = f_n(\lambda/\lambda_1)/\lambda_1$, which converges to a stable distribution as $n$ increases and exhibits a power-law scaling for small $\lambda$ (Figure 3c in the main text):

$$f_n(x) \sim x^{-\alpha}, \tag{S35}$$

with a lower cutoff, $x_{\min}(n) \sim n^{-\beta}$ (Figure 3e in the main text).

We remark that the scaling expression of the added viscosity due to crosslinks (Eq. (S30)) implies a constraint on the distribution of small eigenvalues. Clearly, $\Delta\eta$ must be dominated by the eigenvalues of the slow modes between 0 and $\lambda_1$ (recall that $p_n(\lambda) \sim \lambda^{-1/3}$ for small $\lambda$), which leads to

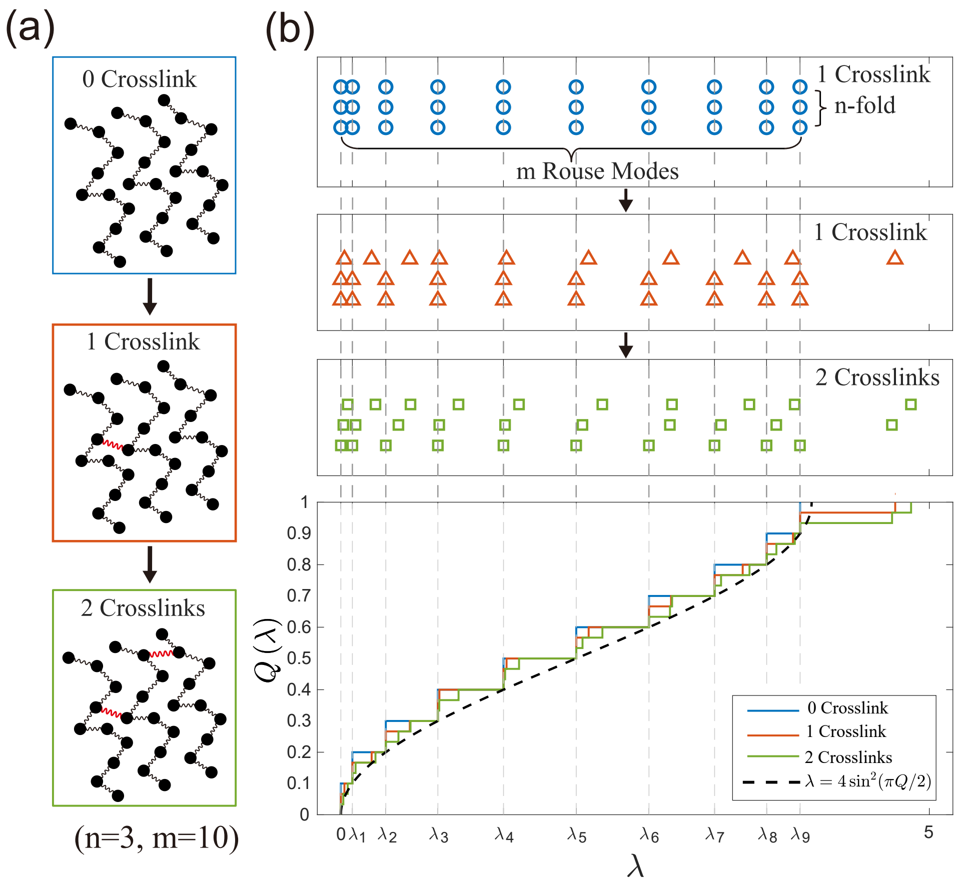


FIG. S2. Eigenvalues for a single cluster. (a) A schematic of how a 3-chain cluster is constructed by adding crosslink one by one. In this example, each chain has $m = 10$ beads. (b) Visualization of the eigenvalues as the number of crosslinks increases, corresponding to (a). The vertical dashed lines mark the $m$ distinct Rouse modes ($\lambda_0$ through $\lambda_9$) of the uncrosslinked chains. The upper panels display the discrete eigenvalues for the initial uncrosslinked chains (blue), a single-crosslink intermediate (red), and the final cluster (green). The bottom panel shows the evolution of the cumulative distribution of the eigenvalues, $Q(\lambda)$. The black dashed curve is the analytical relationship $\lambda = 4\sin^2(\pi Q/2)$ from Eq. (S34), which holds exactly for the discrete Rouse modes where $Q(\lambda_q) = q/m$.

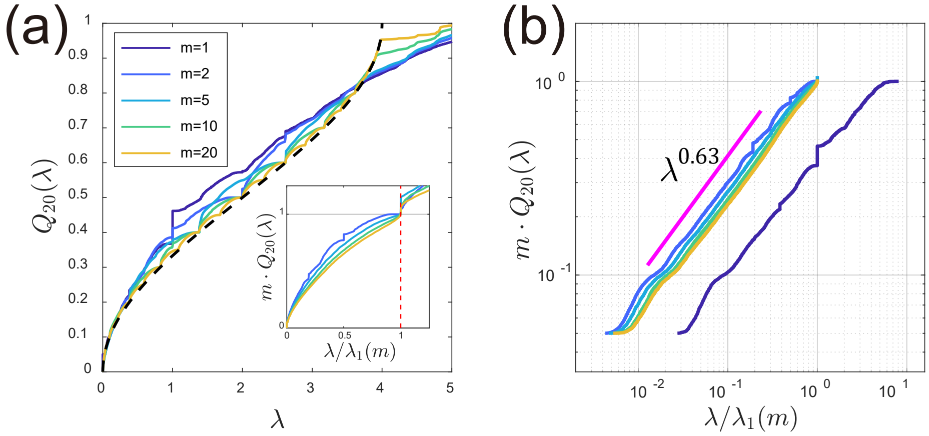


FIG. S3. (a) The cumulative eigenvalue distributions for clusters of size $n = 20$ under different values of $m$. The inset shows the distribution of eigenvalues for $m \geq 2$ between 0 and $\lambda_1$ after normalization by $\lambda_1$, which converges to a limit for sufficiently large $m$. (b) The same as the inset of (a), but in a log-log plot. Here, the $m = 1$ curve is also shown by taking $\lambda_1(m = 1) = 1$. We note that the scaling of the eigenvalue distributions in the small $\lambda$ limit is independent of $m$.

$$\begin{aligned}\Delta\eta &\approx \sum_{0<\lambda<\lambda_1} \frac{1}{\lambda} \approx nm \int_{\lambda_{\min}}^{\lambda_1} \frac{1}{\lambda} p_n(\lambda) d\lambda \\ &\sim nm \int_{\lambda_{\min}}^{\lambda_1} \frac{1}{\lambda}\frac{1}{m\lambda_1}\left(\frac{\lambda}{\lambda_1}\right)^{-\alpha} d\lambda \\ &\sim n\lambda_1^{\alpha-1}\int_{\lambda_{\min}}^{\lambda_1} \lambda^{-(\alpha+1)} d\lambda \\ &\sim n\lambda_1^{\alpha-1}\lambda_{\min}^{-\alpha} \\ &\sim m^2 n^{1+\alpha\beta}.\end{aligned} \tag{S36}$$

Comparing Eq. (S30) with Eq. (S36), we predict that

$$\alpha\beta = \frac{1}{2}, \tag{S37}$$

which agrees perfectly with our numerical calculations where we find $\alpha \approx 1/3$ and $\beta \approx 3/2$ (Figure 3c and e in the main text).

### D. Properties of an ER Random Graph

#### *1. Physical Justification and Derivation of the Branch Size Distribution*

Before deriving the analytical branch size distribution, we first provide the physical justification for mapping the condensate's network topology to an Erdős-Rényi (ER) random graph. Classical theories of polymer physics have well established that polymer chains strongly overlap within the dense phase of a phase-separated polymer solution (i.e., the condensate), and the screening effects ensure that the chain configuration adopts Gaussian statistics [6], in agreement with recent computational simulations of biomolecular condensates [11–13]. Therefore, the overlap parameter—defined as the average number of other chains within the pervaded volume of a given chain—is much larger than one. Consequently, for a cluster of crosslinked chains seeking to form a new crosslink, the number of available reactive sites on other chains vastly exceeds those within the cluster itself. This statistical imbalance makes the formation of loops (including multiple crosslinks between the same two chains or intramolecular crosslinks) essentially negligible.

Furthermore, because we focus on the liquid-like state where the mean number of crosslinks per chain $c < 1$, any cluster growth process is guaranteed to terminate, resulting exclusively in finite-sized clusters. Mathematically, if we pick a random chain and explore its connected component, this process is identical to a Galton-Watson branching process [14]. Starting from a single chain (the "progenitor"), the cluster grows by adding new offspring, whose number follows a Poisson distribution with mean $c$. Because of the absence of loops and the constant crosslink probability, the network structure of the entire condensate seamlessly maps to a subcritical ER random graph $G(N_p, c)$ in the thermodynamic limit (where the number of chains $N_p \to \infty$). In this configuration, the graph decomposes into an ensemble of finite, tree-like clusters without loops. Therefore, the degree distribution of a randomly chosen node (chain) follows a Poisson distribution:

$$P(\text{deg} = k_1) = \frac{e^{-c}c^{k_1}}{k_1!}. \tag{S38}$$

If we randomly select an edge and randomly choose one of its connected nodes, the distribution of the remaining degree $k_2$ (i.e., the number of edges incident to the node excluding the initially selected edge) is:

$$P(\text{remaining deg} = k_2) = \frac{P(\text{deg} = k_2+1)(k_2+1)}{\sum_k P(\text{deg} = k)k}. \tag{S39}$$

Since $\sum_k P(k)k = c$, this simplifies to:

$$P(\text{remaining deg} = k_2) = \frac{e^{-c}c^{k_2}}{k_2!}, \tag{S40}$$

which is identical to the original Poisson distribution. We introduce the generating function for Eq. (S38):

$$G_p(x) = \sum_k P(k)x^k = e^{c(x-1)}. \tag{S41}$$

To analyze branch sizes, we define a branch selection process: (1) randomly select an edge, (2) cut it to split the component into two branches, and (3) choose one branch randomly. Let $P(\text{size} = n)$ be the size distribution of the selected branch and $G(x)$ be its generating function. Starting from the root node of the chosen branch, let $k_2$ denote its remaining degree (excluding the cut edge). The root connects to $k_2$ sub-branches with sizes $n_1, n_2, \ldots, n_{k_2}$, giving total branch size $n = 1 + \sum_{i=1}^{k_2} n_i$. For fixed $k_2$, the conditional generating function is $G(x|k_2) = xG(x)^{k_2}$. Averaging over all possible $k_2$ and using the generating function $G_p(z) = e^{c(z-1)}$ of the remaining degree distribution, we obtain the recursive relation:

$$G(x) = \mathbb{E}_{k_2}[xG(x)^{k_2}] = xG_p(G(x)) = xe^{c(G(x)-1)}. \tag{S42}$$

We set $u = G(x)$, so $x(u) = ue^{c(1-u)}$. The Lagrange inversion formula gives the coefficient of $x^n$ in $G(x)$ [15]:

$$[x^n]G(x) = \frac{1}{n}[u^{n-1}]\left(\frac{u}{x(u)}\right)^n. \tag{S43}$$

Therefore,

$$[x^n]G(x) = \frac{1}{n}[u^{n-1}]\left(\frac{u}{ue^{c(1-u)}}\right)^n = \frac{1}{n}[u^{n-1}]e^{nc(u-1)}. \tag{S44}$$

We expand the exponential:

$$e^{ncu} = \sum_{k=0}^{\infty} \frac{(ncu)^k}{k!} \tag{S45}$$

so the coefficient of $u^{n-1}$ in $e^{ncu}$ is $\frac{(nc)^{n-1}}{(n-1)!}$. Therefore,

$$[u^{n-1}]e^{ncu} = \frac{(nc)^{n-1}}{(n-1)!} \tag{S46}$$

and

$$[x^n]G(x) = \frac{1}{n}e^{-nc}\frac{(nc)^{n-1}}{(n-1)!} = \frac{e^{-nc}(nc)^{n-1}}{n!}. \tag{S47}$$

Thus, the branch size distribution is:

$$P(n) = \frac{e^{-nc}(nc)^{n-1}}{n!}. \tag{S48}$$

The asymptotic behavior for large $n$ is found using Stirling's approximation $n! \approx \sqrt{2\pi n}(n/e)^n$:

$$P(n) \approx \frac{e^{-nc}(nc)^{n-1}}{\sqrt{2\pi n}(n/e)^n} = \frac{c^{n-1}e^{n(1-c)}}{\sqrt{2\pi}n^{3/2}}. \tag{S49}$$

Finally, we remark that the branch size distribution, Eq. (S48), is identical to the cluster size distribution we introduce in the main text, i.e., the probability distribution for a randomly selected chain to belong to a cluster of size $n$. This is because the generating function of the cluster size distribution $G'(x)$ can also be written as $G'(x) = xe^{c(G(x)-1)}$, which means that $G'(x) = G(x)$.

#### *2. Viscosity of an ER Random Graph*

Using Eq. (S22), we compute the dimensionless viscosity of the coarse-grained polymer solution where each linear chain is coarse-grained as a node. We sum over all edges, with each edge's contribution equal to the number of paths traversing it:

$$\begin{aligned}\eta_{ER} &= \frac{N_p c}{2}\sum_{n_1=1}^{\infty}\sum_{n_2=1}^{\infty} P(n_1)P(n_2)\frac{n_1 n_2}{n_1+n_2} \\ &= \sum_{N\geq 2}\sum_{n=1}^{N-1}\frac{N_p e^{-cN}c^{N-1}n^n(N-n)^{N-n}}{2Nn!(N-n)!} \\ &= \sum_{N\geq 2}\eta_N(c).\end{aligned} \tag{S50}$$

Here, $N_p$ is the total number of nodes, $P(n) = e^{-nc}(nc)^{n-1}/n!$ is the distribution of branch size by randomly choosing an edge, and $\eta_N(c)$ is the viscosity contribution of all clusters of size $N$. Applying Stirling's approximation $n^n/n! \approx e^n/\sqrt{2\pi n}$ to Eq. (S50) yields

$$\begin{aligned}\eta_N(c)/N_p &\approx \frac{e^{(1-c)N}c^{N-1}}{4\pi N}\sum_{n_1=1}^{N-1}\frac{1}{\sqrt{n_1(N-n_1)}} \\ &\approx \frac{e^{(1-c)N}c^{N-1}}{4\pi N}\int_0^1\frac{dx}{\sqrt{x(1-x)}} \\ &\approx \frac{e^{(1-c+\ln c)N}}{4cN}.\end{aligned} \tag{S51}$$

In the weakly crosslinked limit ($c \ll 1$), the dimensionless viscosity of the coarse-grained network is dominated by the smallest cluster size ($N = 2$), yielding $\eta_{ER} = N_p c/4$ using Eq. (S50). As expected, the added viscosity is proportional to $c$ in the weakly crosslinked limit. Near the gelation point ($\epsilon = 1 - c \ll 1$), $\eta_N(c)$ can be further approximated as $\eta_N(c) \approx N_p e^{-\epsilon^2 N/2}/4N$, from which we find a logarithmic divergence of the dimensionless viscosity of the coarse-grained network near the gelation point: $\eta_{ER} \approx N_p \sum_{N\geq 2} e^{-\epsilon^2 N/2}/4N \approx -(N_p/2)\ln\epsilon$. Using the exact formula, Eq. (S50), we compute the viscosity of the coarse-grained ER random graph for $0 < c < 1$ (Figure S4).

It is noteworthy that near the gelation point, the number fraction of cluster size for the ER random graph follows a power law $P'(N) \propto N^{-5/2}$, and each cluster exhibits a fractal dimension of 4 (i.e., $\langle R_g^2\rangle \propto N^{1/2}$) [6]. According to Eq. (S20), the total viscosity of clusters of size $N$ is: $\eta_N \propto P'(N)\cdot N\langle R_g^2\rangle \sim N^{-1}$. This summation elucidates the logarithmic divergence of viscosity near the gelation point, i.e., $\eta_{ER} \sim -\ln\epsilon$. The scaling behavior reveals a universal feature wherein clusters across all scales contribute comparably to the divergence, as the integrand $N^{-1}$ maintains comparable weight for different cluster sizes.

We summarize the behavior of the viscosity for the entire condensate. For long precursor chains ($m \gg 1$), the viscosity increment scales as $\Delta\eta \sim \frac{m^2}{3}\eta_{ER}$, where $\eta_{ER}$ is the viscosity of the coarse-grained network. As discussed, $\eta_{ER}$ varies with the crosslinking density $c$: in the weakly crosslinked limit ($c \ll 1$), $\eta_{ER} \sim c$, leading to $\Delta\eta \propto m^2 N_p c$; near the gelation point ($c \to 1$), $\eta_{ER} \sim -\ln(1-c)$, resulting in $\Delta\eta \propto m^2 N_p(-\ln(1-c))$. This demonstrates how the viscosity evolves with crosslinking density across the entire range below the gelation point (Figure S4).

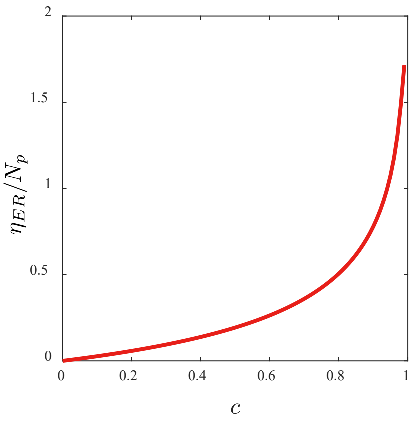


FIG. S4. The viscosity divided by the total number of nodes $\eta_{ER}/N_p$ as a function of crosslinking density $c$ for the coarse-grained ER graph corresponding to a random crosslinked polymer solution. The viscosity increases with crosslinking density, rising rapidly as $c$ approaches the gelation point at $c = 1$.

#### 3. Cluster Topology Distribution of Given Size and Its Sampling Procedure

A crucial assumption in our model is that the full eigenvalue distribution for the condensate can be obtained by averaging the eigenvalue distributions for clusters of size $n$, weighted by the cluster size distribution $P(n)$. This relies on the principle that the relative probability of observing any specific cluster topology of a given size $n$ is independent of the mean degree $c$.

To demonstrate the validity of this assumption, we consider an Erdős-Rényi (ER) random graph model $G(N_p, p_{\text{edge}})$, where $N_p$ is the total number of polymer chains (nodes) and $p_{\text{edge}}$ is the probability of an edge (crosslink) existing between any two nodes. In the thermodynamic limit, this is equivalent to our model with $p_{\text{edge}} = c/(N_p - 1)$. Consider a specific set of $n$ nodes. For this set to form a specific, isolated, labeled cluster $T$ of size $n$ (see examples of labeled clusters in Figure S5), two conditions must be met. First, the specific $n-1$ edges that define the topology of $T$ must be present, the probability for which is $p_{\text{edge}}^{n-1}$. Second, all other potential edges that would change the cluster's structure or connect the cluster to other nodes must be absent. The probability of the latter is $(1 - p_{\text{edge}})^{\binom{n}{2} - (n-1) + n(N_p - n)}$. To summarize, the probability of forming the specific, isolated, labeled cluster $T$ is the product of these terms. We note that this probability depends only on $n$ and $p_{\text{edge}}$, but not on the specific topology of the cluster $T$, since every cluster of size $n$ has exactly $n-1$ edges in our case. Consequently, any other labeled cluster $T'$ of the same size has the same probability to form as an isolated cluster. Therefore, the conditional probability of observing a specific topology for all clusters of size $n$ should be independent of $p_{\text{edge}}$ and $c$.

To ensure our sampling of clusters is free from finite-size effects, we follow a two-step procedure. First, we generate the coarse-grained topology of a random, isolated cluster of size $n$ by directly generating a random labeled cluster using Prüfer codes [16]. By Cayley's formula, there is a one-to-one correspondence between the $n^{n-2}$ distinct labeled clusters on $n$ nodes and codes of length $n-2$. An illustration of this correspondence is provided in Figure S5. We thus construct a random Prüfer code and deterministically convert it to its unique corresponding cluster. This guarantees an unbiased sampling over all possible labeled topologies. We highlight that this sampling is designed to be statistically identical to drawing all clusters of size $n$ from an ER graph in the thermodynamic limit.

Second, we expand this $n$-node graph into a full cluster of $n \times m$ beads. Each of the $n$ nodes from the initial step is restored into a linear Rouse chain composed of $m$ beads. The $n-1$ edges of the coarse-grained topology dictate the intermolecular crosslinks: for each such edge connecting two chains, a crosslink is formed by adding an edge between one randomly selected bead from each of the two chains. This process results in a cluster of $n \times m$ total beads. We then construct the graph Laplacian for this final network and numerically compute its entire eigenvalue distribution.

To compute the eigenvalue distribution for clusters of fixed size averaged over all possible topologies (Figure 3a, c in the main text), we generate a large ensemble of clusters of a given size $n$ with random labeled topologies, where each chain is labeled by a number (Figure S5). The eigenvalue distribution is then obtained by averaging over all sampled topologies.

### E. Derivation of Dynamic Moduli

The frequency-dependent storage modulus, $G'(\omega)$, and loss modulus, $G''(\omega)$, are the real and imaginary parts of the complex modulus, $G^*(\omega) = G'(\omega) + iG''(\omega)$. The complex modulus is related to the relaxation modulus $G(t)$ via [5, 6]

$$G^*(\omega) = i\omega \int_0^\infty G(t) e^{-i\omega t} dt,$$

from which we obtain that

$$G'(\omega) = \omega \int_0^\infty G(t) \sin(\omega t) dt, \tag{S52}$$

$$G''(\omega) = \omega \int_0^\infty G(t) \cos(\omega t) dt. \tag{S53}$$

For the generalized Rouse model with discrete eigenvalues, the dimensionless relaxation modulus can be written as

$$G(t) = \sum_{\lambda \neq 0} e^{-\lambda t},$$

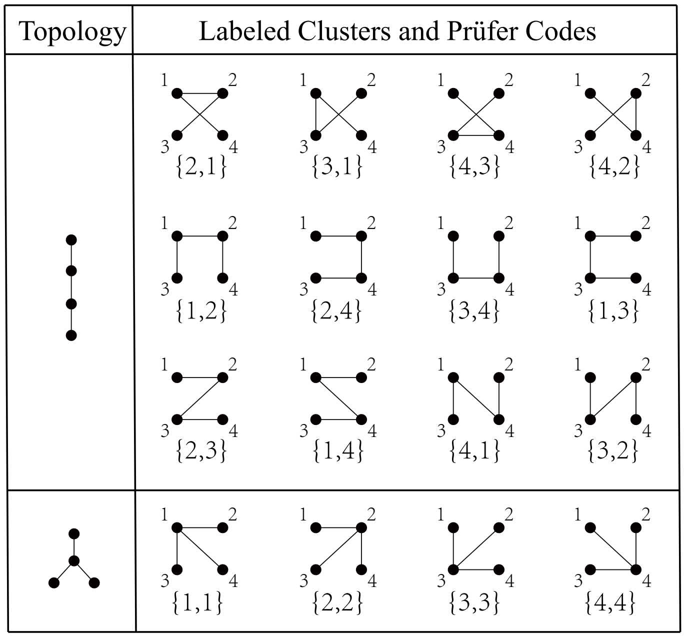


FIG. S5. Illustration of the one-to-one correspondence between labeled clusters and Prüfer codes for $n = 4$ nodes. Two distinct topologies are shown, each associated with its unique Prüfer code of length $n - 2 = 2$.

for $t > 0$. Substituting this expression into the definition of $G^*(\omega)$, we find that

$$G^*(\omega) = \sum_{\lambda \neq 0} i\omega \int_0^\infty e^{-(\lambda + i\omega)t} dt = \sum_{\lambda \neq 0} \frac{i\omega}{\lambda + i\omega} = \sum_{\lambda \neq 0} \frac{\omega^2 + i\omega\lambda}{\lambda^2 + \omega^2},$$

from which we find the storage and loss moduli as

$$G'(\omega) = \sum_{\lambda \neq 0} \frac{\omega^2}{\lambda^2 + \omega^2}, \tag{S54}$$

$$G''(\omega) = \sum_{\lambda \neq 0} \frac{\omega\lambda}{\lambda^2 + \omega^2}. \tag{S55}$$

To calculate the storage modulus $G'(\omega)$ and loss modulus $G''(\omega)$, we recall the scaling behavior of the relaxation modulus $G(t)$ as established in the main text,

$$G(t) \sim \begin{cases} N_p m & t \ll 1 \\ \frac{N_p m}{\sqrt{t}} & 1 \ll t \ll \lambda_1^{-1} \\ \frac{N_p}{\lambda_1 t} & \lambda_1^{-1} \ll t \ll \lambda_{\min}^{-1} \\ \frac{N_p}{\lambda_1 t} e^{-\lambda_{\min} t} & t \gg \lambda_{\min}^{-1}. \end{cases} \tag{S56}$$

In writing the above equations, we have used the condition that $m \gg 1$ so that $\lambda_1 \sim (1/m^2) \ll 1$ and $c \to 1$ so that $\lambda_{\min}/\lambda_1 \sim (1-c)^{2\beta} \ll 1$.

To proceed rigorously, we first justify the scaling $G(t) \sim N_p m/\sqrt{t}$ for the crosslinked system in the time regime $1 \ll t \ll \lambda_1^{-1}$. The relaxation modulus of the entire condensate, $G(t) = \sum_{\lambda \neq 0} e^{-\lambda t}$, is a sum over the non-zero eigenvalues of the condensate's connectivity matrix formed from a system of $N_p$ initially disconnected Rouse chains (each with $m$ beads) by the addition of $N_p c/2$ crosslinks. In the uncrosslinked state, there is an $N_p$-fold degeneracy of each Rouse mode $\lambda_q$ ($q = 0, 1, \cdots, m-1$). According to the eigenvalue interlacing theorem as demonstrated in Section

C and Eq. (S31), after the addition of $N_pc/2$ crosslinks, $N_pc/2$ eigenvalues are placed in each interval $(\lambda_q, \lambda_{q+1})$ for $q = 0, 1, \cdots, m-2$ and $(\lambda_{m-1}, \infty)$, and $N_p(1-c/2)$ eigenvalues remain at $\lambda_q$.

A lower bound, $G_{\text{lower}}(t)$, is obtained by increasing each eigenvalue. Specifically, we increase the non-zero eigenvalues in $(0, \lambda_1)$ and those in $[\lambda_{m-1}, \infty)$ to infinity. For $q = 1, \ldots, m-2$, we increase the $N_p$ eigenvalues in $[\lambda_q, \lambda_{q+1})$ to $\lambda_{q+1}$. This yields:

$$G_{\text{lower}}(t) = N_p \sum_{k=2}^{m-1} e^{-\lambda_k t}. \tag{S57}$$

Conversely, an upper bound, $G_{\text{upper}}(t)$, is obtained by decreasing each eigenvalue. We decrease the $N_pc/2$ non-zero eigenvalues in $(0, \lambda_1)$ to zero. For the intervals $q \geq 1$ (including $[\lambda_{m-1}, \infty)$), the $N_p$ eigenvalues inside $[\lambda_q, \lambda_{q+1})$ are decreased to $\lambda_q$. This gives:

$$G_{\text{upper}}(t) = N_pc/2 + N_p \sum_{k=1}^{m-1} e^{-\lambda_k t} = N_pc/2 + G_{\text{Rouse}}(t), \tag{S58}$$

where $G_{\text{Rouse}}(t) = N_p \sum_{k=1}^{m-1} e^{-\lambda_k t} \sim N_p m \int_{\lambda_1}^{1} \lambda^{-1/2} e^{-\lambda t} d\lambda \sim N_p m/\sqrt{t}$ $(1 \ll t \ll \lambda_1^{-1})$ is the uncrosslinked modulus.

Notably, both $G(t)$ and the uncrosslinked modulus $G_{\text{Rouse}}(t)$ are in the interval $(G_{\text{lower}}(t), G_{\text{upper}}(t))$. Therefore, $|G(t) - G_{\text{Rouse}}(t)| < G_{\text{upper}}(t) - G_{\text{lower}}(t) = N_pc/2 + N_p e^{-\lambda_1 t} < 2N_p$. The relative error between $G(t)$ and $G_{\text{Rouse}}(t)$ is therefore limited by:

$$\frac{|G(t) - G_{\text{Rouse}}(t)|}{G_{\text{Rouse}}(t)} < \frac{2N_p}{N_p m/\sqrt{t}}. \tag{S59}$$

Given the time regime $1 \ll t \ll \lambda_1^{-1} \sim m^2$, this relative error is negligible, justifying the approximation of $G(t)$ with the Rouse scaling.

As for $t \ll 1$, the vast majority of the eigenvalues are below 4 since $N_p(m-1)$ eigenvalues are smaller than $\lambda_{m-1}$, which is smaller than 4. Consequently, for almost all modes, $\lambda t < 4t \ll 1$, leading to $e^{-\lambda t} \approx 1$. Summing over these modes yields the plateau modulus $G(t) \approx N_p m$.

In the following, we calculate the storage modulus $G'(\omega)$ and loss modulus $G''(\omega)$ for different frequency regimes.

#### 1. *High-Frequency Limit ($\omega \gg 1$)*

In the limit where the frequency is much larger than any eigenvalues ($\omega \gg 1$; note that the largest eigenvalue $\lambda_{\max} \sim 1$), the storage and loss moduli can be directly found via Eq. (S55) as

$$G'(\omega) = \sum_{\lambda \neq 0} \frac{\omega^2}{\lambda^2 + \omega^2} \approx \sum_{\lambda \neq 0} \frac{\omega^2}{\omega^2} = N_p m, \tag{S60}$$

$$G''(\omega) = \sum_{\lambda \neq 0} \frac{\omega \lambda}{\lambda^2 + \omega^2} \approx \sum_{\lambda \neq 0} \frac{\omega \lambda}{\omega^2} = \frac{N_p m}{\omega}. \tag{S61}$$

Here, we have used the fact that there are $N_p m$ eigenvalues in total.

#### 2. *Rouse Regime ($\lambda_1 \ll \omega \ll 1$)*

In this regime, corresponding to short times $(1 \ll t \ll \lambda_1^{-1})$, the relaxation modulus is dominated by the Rouse modes and scales as $G(t) \sim N_p m/\sqrt{t}$. We first calculate the Fourier transform of $t^{-1/2}$:

$$\int_0^\infty t^{-1/2} e^{-i\omega t} dt = \sqrt{\frac{\pi}{\omega}} e^{-i\pi/4} = \sqrt{\frac{\pi}{2\omega}} (1 - i) \tag{S62}$$

Substituting this into the definition of the complex modulus $G^*(\omega) = i\omega \int G(t) e^{-i\omega t} dt$:

$$G^*(\omega) \sim i\omega \left( N_p m \sqrt{\frac{\pi}{2\omega}}(1-i) \right) = N_p m \sqrt{\frac{\pi\omega}{2}}(i+1), \tag{S63}$$

from which we find the storage and loss moduli as

$$G'(\omega) \sim N_p m \sqrt{\omega}, \tag{S64}$$
$$G''(\omega) \sim N_p m \sqrt{\omega}, \tag{S65}$$

which is the characteristic $\sqrt{\omega}$ scaling of the Rouse model.

#### *3. Collective-Behavior Regime ($\lambda_{\min} \ll \omega \ll \lambda_1$)*

This regime corresponds to $\lambda_1^{-1} \ll t \ll \lambda_{\min}^{-1}$, where the modulus scales as $G(t) \sim N_p/(\lambda_1 t)$, from which we find

$$G'(\omega) \sim \omega \int_{\lambda_1^{-1}}^{\lambda_{\min}^{-1}} \frac{N_p}{\lambda_1 t} \sin(\omega t) dt \sim N_p(\omega/\lambda_1)[Si(\omega/\lambda_{\min}) - Si(\omega/\lambda_1)], \tag{S66}$$

$$G''(\omega) \sim \omega \int_{\lambda_1^{-1}}^{\lambda_{\min}^{-1}} \frac{N_p}{\lambda_1 t} \cos(\omega t) dt \sim N_p(\omega/\lambda_1)[Ci(\omega/\lambda_{\min}) - Ci(\omega/\lambda_1)], \tag{S67}$$

where we introduce the Sine Integral ($\mathrm{Si}(x)$) and Cosine Integral ($\mathrm{Ci}(x)$) functions,

$$\mathrm{Si}(x) = \int_0^x \frac{\sin t}{t} dt, \tag{S68}$$

$$\mathrm{Ci}(x) = -\int_x^\infty \frac{\cos t}{t} dt. \tag{S69}$$

In the regime where $\lambda_{\min} \ll \omega \ll \lambda_1$, we have $\omega/\lambda_1 \ll 1$ and $\omega/\lambda_{\min} \gg 1$. Thus, we can use the asymptotic behaviors: $\mathrm{Si}(x \to 0) \to 0$, $\mathrm{Si}(x \to \infty) \to \pi/2$, $\mathrm{Ci}(x \to 0) \to \ln(x)$, and $\mathrm{Ci}(x \to \infty) \to 0$. We find the storage modulus as

$$G'(\omega) \sim N_p(\omega/\lambda_1)\left(\mathrm{Si}(\omega/\lambda_{\min}) - \mathrm{Si}(\omega/\lambda_1)\right) \approx N_p(\omega/\lambda_1)\left(\frac{\pi}{2} - 0\right) \sim N_p(\omega/\lambda_1), \tag{S70}$$

and the loss modulus as

$$G''(\omega) \sim N_p(\omega/\lambda_1)\left(\mathrm{Ci}(\omega/\lambda_{\min}) - \mathrm{Ci}(\omega/\lambda_1)\right) \approx N_p(\omega/\lambda_1)\left(0 - \ln\left(\frac{\omega}{\lambda_1}\right)\right) = N_p(\omega/\lambda_1)\ln\left(\frac{\lambda_1}{\omega}\right). \tag{S71}$$

#### *4. Low-Frequency Limit ($\omega \ll \lambda_{\min}$)*

This regime is sensitive to the behavior of $G(t)$ at long times; for $t \gg \lambda_{\min}^{-1}$, the decay is approximately exponential, $G(t) \sim N_p e^{-\lambda_{\min} t}/(\lambda_1 t)$. Since $\omega$ is very small in this regime, we can make approximations such that $\sin(\omega t) \approx \omega t$ and $\cos(\omega t) \approx 1$ as long as $t < \lambda_{\min}^{-1}$. We find the storage modulus $G'(\omega)$ as

$$\begin{aligned}
G'(\omega) &\sim \omega \left( \int_{\lambda_1^{-1}}^{\lambda_{\min}^{-1}} \frac{N_p}{\lambda_1 t} \sin(\omega t) dt + \int_{\lambda_{\min}^{-1}}^{\infty} \frac{N_p e^{-\lambda_{\min} t}}{\lambda_1 t} \sin(\omega t) dt \right) \\
&\approx N_p(\omega/\lambda_1) \left( \int_{\lambda_1^{-1}}^{\lambda_{\min}^{-1}} \frac{\omega t}{t} dt + \int_{\lambda_{\min}^{-1}}^{\infty} \frac{e^{-\lambda_{\min} t}(\omega t)}{t} dt \right) \\
&\approx N_p(\omega^2/\lambda_1) \left( \left( \frac{1}{\lambda_{\min}} - \frac{1}{\lambda_1} \right) + \int_{\lambda_{\min}^{-1}}^{\infty} e^{-\lambda_{\min} t} dt \right) \\
&\approx N_p(\omega^2/\lambda_1) \left( \frac{1}{\lambda_{\min}} - \frac{1}{\lambda_1} + \frac{1}{\lambda_{\min}} e^{-1} \right) \\
&\sim N_p \omega^2/(\lambda_1 \lambda_{\min}) \\
&\sim N_p(\omega/\lambda_1)^2 (1-c)^{-2\beta},
\end{aligned} \tag{S72}$$

and the loss modulus $G''(\omega)$ as

$$
\begin{aligned}
G''(\omega) &\sim \omega\left(\int_{\lambda_1^{-1}}^{\lambda_{\min}^{-1}} \frac{N_p}{\lambda_1 t}\cos(\omega t)dt + \int_{\lambda_{\min}^{-1}}^{\infty} \frac{N_p e^{-\lambda_{\min} t}}{\lambda_1 t}\cos(\omega t)dt\right) \\
&\approx N_p(\omega/\lambda_1)\left(\int_{\lambda_1^{-1}}^{\lambda_{\min}^{-1}} \frac{1}{t}dt + \int_{\lambda_{\min}^{-1}}^{\infty} \frac{e^{-\lambda_{\min} t}}{t}dt\right) \\
&\approx N_p(\omega/\lambda_1)\left(\ln\left(\frac{\lambda_1}{\lambda_{\min}}\right) + E_1\right),
\end{aligned}
\tag{S73}
$$

where $E_1 = \int_1^\infty (e^{-x}/x)dx$, which is a constant. The leading term is linear in frequency, giving the scaling $G''(\omega) \sim N_p(\omega/\lambda_1)\ln(\lambda_1/\lambda_{\min}) \sim -N_p m^2 \ln(1-c)\omega$. In summary, combining the results from all four regimes gives the following scaling behaviors:

$$
G'(\omega) \sim \begin{cases} N_p\omega^2/(\lambda_1\lambda_{\min}) & \omega \ll \lambda_{\min} \\ N_p\omega/\lambda_1 & \lambda_{\min} \ll \omega \ll \lambda_1 \\ N_p m\sqrt{\omega} & \lambda_1 \ll \omega \ll 1 \\ N_p m & \omega \gg 1, \end{cases}
\tag{S74}
$$

$$
G''(\omega) \sim \begin{cases} N_p(\omega/\lambda_1)\ln(\frac{\lambda_1}{\lambda_{\min}}) & \omega \ll \lambda_{\min} \\ N_p(\omega/\lambda_1)\ln(\frac{\lambda_1}{\omega}) & \lambda_{\min} \ll \omega \ll \lambda_1 \\ N_p m\sqrt{\omega} & \lambda_1 \ll \omega \ll 1 \\ N_p m/\omega & \omega \gg 1. \end{cases}
\tag{S75}
$$

We also verify that the universal scaling form of dynamic moduli at low frequencies is independent of $m$ by normalizing $\omega$ with $\lambda_1$ (Eqs. (S74, S75); Figure S6).

#### F. Parameter Selection and Experimental Fitting

We note that the total number of polymer chains, $N_p$, does not affect the fit to the experimental data, as it cancels out in the normalized moduli. Meanwhile, the number of beads per chain $m$ sets the width of the intermediate $G'(\omega) \sim \omega^{1/2}$ scaling regime by setting its lower frequency bound, $\lambda_1$ (Figure S6). The predicted curves with $m = 5$ reasonably match the experimental data, whereas those with larger $m$ deviate in the high-frequency regime (Figure S7b). We notice that some data of Ref. [17] are from condensates containing a pentameric protein with five folded domains joined by flexible linkers, consistent with the fitting parameter $m = 5$ according to our model. Regarding the average number of crosslinks per chain $c$, it determines the transition between the linear-scaling regime ($G' \sim \omega$) and the Maxwell-fluid regime ($G' \sim \omega^2$). As the system approaches the gelation point, the crossover shifts to extremely low frequencies, which can be beyond the reach of experimental measurements. We note that the predicted curves with $c = 0.8$ also fit the experimental data reasonably well except for the deviation at low frequencies (Figure S7c).

[1] Prince E Rouse. A theory of the linear viscoelastic properties of dilute solutions of coiling polymers. *The Journal of Chemical Physics*, 21(7):1272–1280, 1953.
[2] Yuliang Yang. Graph theory of viscoelastic and configurational properties of gaussian chains. *Macromolecular Theory and Simulations*, 7(5):521–549, 1998.

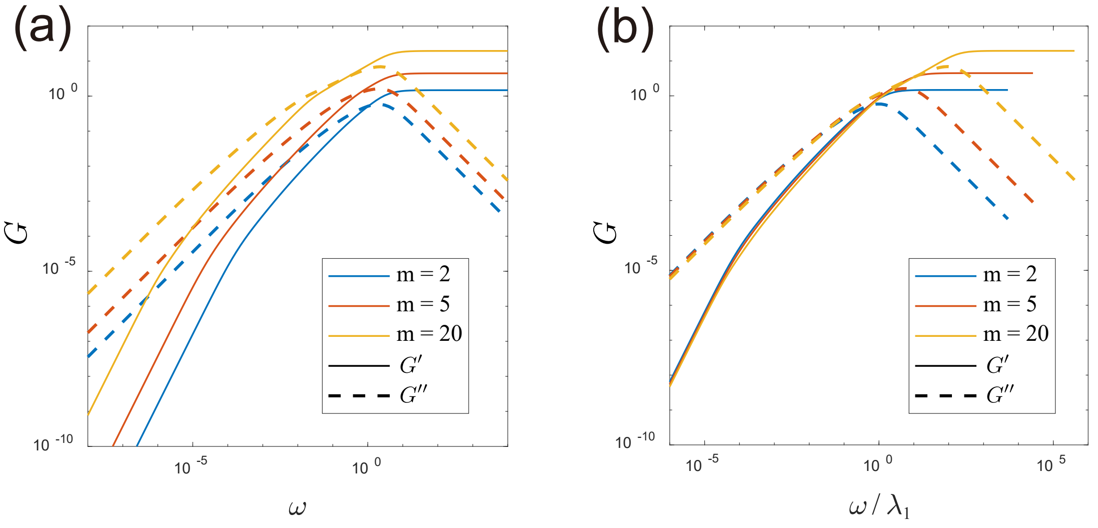


FIG. S6. Universal scaling form of the dynamic moduli for different numbers of beads per chain $m$. (a) The storage modulus $G'$ (solid lines) and loss modulus $G''$ (dashed lines) as a function of the frequency $\omega$ for various numbers of beads per chain $m = 2, 5, 20$ at a fixed mean crosslink number per chain $c = 0.95$. (b) The same moduli plotted against the normalized frequency $\omega/\lambda_1$, where $\lambda_1$ is the first Rouse mode. In the low frequency range ($\omega/\lambda_1 < 1$), the curves for different $m$ collapse onto a single master curve.

[3] Samuel R Cohen, Priya R Banerjee, and Rohit V Pappu. Direct computations of viscoelastic moduli of biomolecular condensates. *The Journal of Chemical Physics*, 161(9):095103, 2024.
[4] Sybren Ruurds De Groot and Peter Mazur. *Non-equilibrium thermodynamics*. Courier Corporation, 2013.
[5] Masao Doi. *Soft matter physics*. Oxford University Press, 2013.
[6] Michael Rubinstein and Ralph H Colby. *Polymer physics*. Oxford university press, 2003.
[7] Lajos Takács. On cayley's formula for counting forests. *Journal of Combinatorial Theory, Series A*, 53(2):321–323, 1990.
[8] Wasin So. Rank one perturbation and its application to the laplacian spectrum of a graph. *Linear and Multilinear Algebra*, 46(3):193–198, 1999.
[9] Essam El Seidy, Salah Eldin Hussein, and Atef Mohamed. Properties of the characteristic polynomials and spectrum of Pn and Cn. *International Journal of Applied Mathematical Research*, 5(2):132–137, 2016.
[10] Masao Doi and Sam F Edwards. *The theory of polymer dynamics*. Oxford University Press, 1988.
[11] Jiahui Wang, Dinesh Sundaravadivelu Devarajan, Keerthivasan Muthukumar, Young C. Kim, Arash Nikoubashman, and Jeetain Mittal. Sequence-dependent conformational transitions of disordered proteins during condensation. *Chem. Sci.*, 15:20056–20063, 2024.
[12] Keerthivasan Muthukumar, Dinesh Sundaravadivelu Devarajan, Young C. Kim, and Jeetain Mittal. Sticky interactions govern sequence-dependent dynamics in biomolecular condensates. *Proceedings of the National Academy of Sciences*, 123(1):e2518384122, 2026.
[13] Gaurav Mitra, Souradeep Ghosh, Kiersten M. Ruff, Ruoyao Zhang, Gaurav Chauhan, and Rohit V. Pappu. Distinguishing near- versus off-critical phase behaviors of intrinsically disordered proteins. *bioRxiv*, 2025.
[14] Henry William Watson and Francis Galton. On the probability of the extinction of families. *The Journal of the Anthropological Institute of Great Britain and Ireland*, 4:138–144, 1875.
[15] Milton Abramowitz and Irene A Stegun. *Handbook of mathematical functions with formulas, graphs, and mathematical tables*, volume 55. US Government printing office, 1948.
[16] Heinz Prüfer. Neuer Beweis eines Satzes über Permutationen. *Archiv der Mathematischen Physik*, 27:742–744, 1918.
[17] Archishman Ghosh, Divya Kota, and Huan-Xiang Zhou. Shear relaxation governs fusion dynamics of biomolecular condensates. *Nature Communications*, 12(1):5995, 2021.

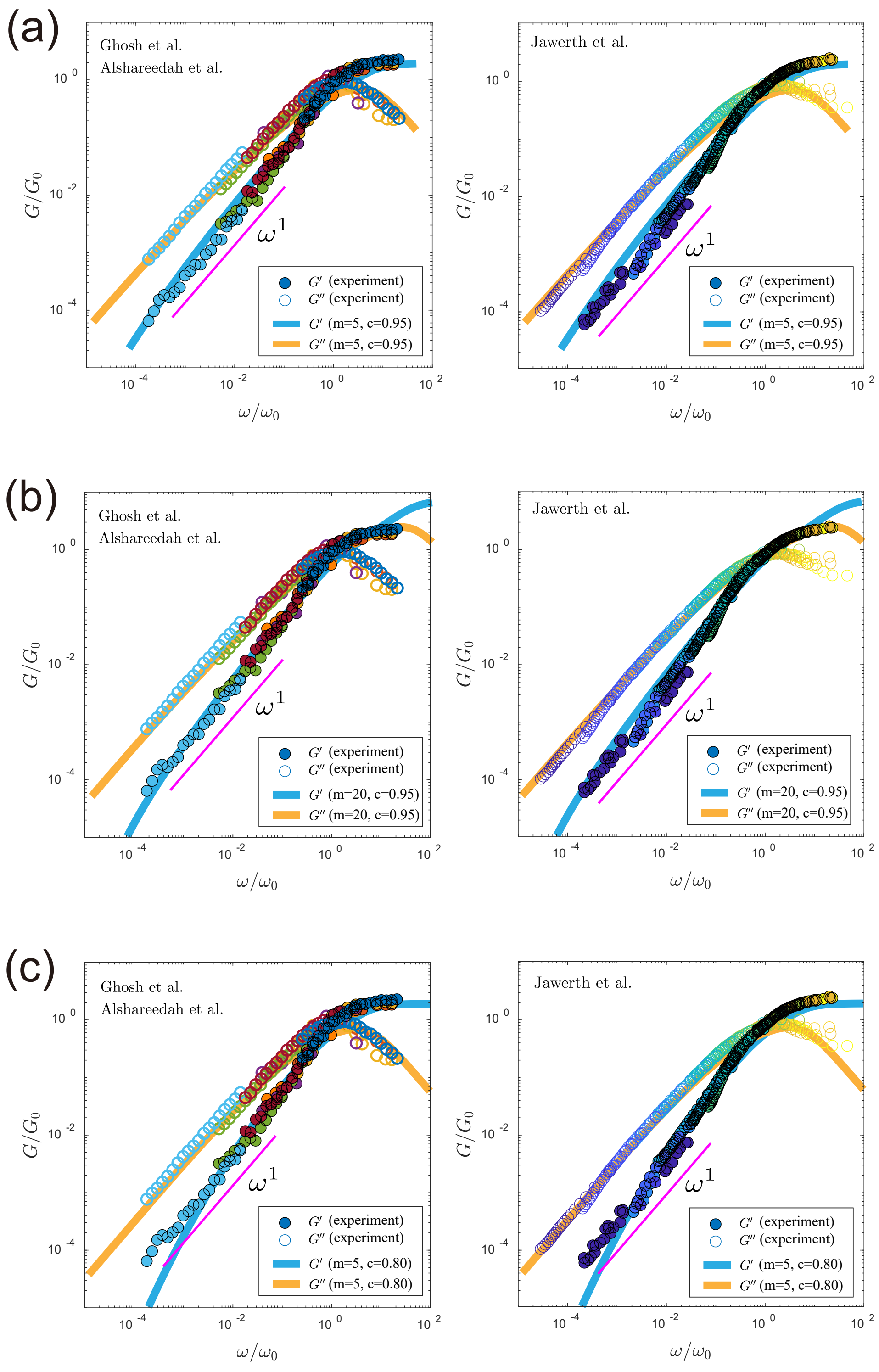


FIG. S7. Comparison of theoretical predictions with experimental data for different parameter sets, demonstrating the distinct constraints on $m$ and $c$. All experimental data are normalized by their respective crossover points $(\omega_0, G_0)$. The solid lines represent the theoretical predictions from our model. (a) The parameters used in the main text ($m = 5$, $c = 0.95$) show good agreement with experimental data. (b) Using a larger number of beads per chain ($m = 20$, $c = 0.95$) results in a deviation at high frequencies. (c) Using a lower crosslink density ($m = 5$, $c = 0.80$) produces a good fit over most of the frequency range, with deviation appearing only at the lowest frequencies. The magenta line has a slope of 1 in the log-log plot.
